# Multiscale reorganization of brain and behavior under large-scale electrical perturbation

**DOI:** 10.64898/2026.03.23.713593

**Authors:** Sarah Kreuzer, Juergen Dukart, Justine Y. Hansen, Hoang K. Nguyen, Michael Bentsch, Sophia Zieger, Katrin Sakreida, Thomas C. Baghai, Caroline Nothdurfter, Michael Grözinger, Bogdan Draganski, Bratislav Mišić, Danilo Bzdok, Simon B. Eickhoff, Timm B. Poeppl

## Abstract

Large-scale electrical perturbation of the human brain provides a unique model for understanding how multiscale biological constraints shape behaviorally relevant reorganization. Here, we integrate longitudinal neuroimaging coordinates from 148 experiments (≈2,300 subjects) with normative connectomics, chemoarchitecture, intrinsic electrophysiology, and transcriptomics to identify cross-scale principles governing human brain reconfiguration under strong perturbation. Convergent hubs of structural and functional plasticity embed within default-mode and salience systems and show complementary coupling to visual networks, linking perturbation-induced change to large-scale circuits supporting affective regulation, memory, interoception, and psychosis-relevant processes. These macroscopic patterns align with intrinsic cortical dynamics and chemoarchitectural gradients dominated by 5-HT_1A_ receptors, with additional contributions from D_2_, μ-opioid and GABA_A_ systems, and are enriched for astrocytic and microglial gene expression, implicating glial plasticity in systems-level reorganization. Finally, in a separate intervention dataset, regularized statistical-learning models demonstrate that this multiscale signature tracks behaviorally relevant symptom change specifically under strong electrical perturbation. Together, these results outline general organizing principles linking molecular, cellular and network-level constraints to human behavioral adaptation, providing a computational framework for understanding how large-scale perturbations reshape brain systems across levels of biological organization.

## INTRODUCTION

In the 1930s, Italian neuropsychiatrist Ugo Cerletti and his assistant Lucio Bini fortuitously noticed that frantic animals in the slaughterhouse were rendered calm by a stunning bolt of electric current to the head, involving a seizure^1,2^. Building on the then-prevalent theories positing an antagonism between schizophrenia and epilepsy – and motivated by Hungarian psychiatrist Ladislaus von Meduna’s poorly controllable and distressing chemically induced convulsions^3^ – Cerletti and Bini replaced pharmacological induction with precisely delivered electrical stimulation. In April 1938, they administered the first electroshock treatment to a psychotic patient using their Bini-Cerletti apparatus, which was later patented^2^. This pivotal experiment laid the foundation for electroconvulsive therapy (ECT) as the most effective intervention for treating well-defined, serious mental illness, including treatment-resistant depression and psychosis. Modern ECT continues to involve the application of electrical current to the brain via externally placed electrodes on the patient’s scalp to induce an epileptic seizure but is nowadays administered under general anesthesia and with muscle relaxation.

Despite decades of clinical application and its phenomenal efficacy with remission rates up to 80%^4^, its mechanism of action remains elusive. Cerletti himself assumed that “acroagonines” – mysterious neuroactive emulsions released by the brain under electroshock – might hold brain-curing properties^2^. More than 80 years later, we still do not fully understand whether it is the applied electricity, or the resulting seizure, or their interaction, that are responsible for the therapeutic effect^5^. Beyond its clinical role, ECT also constitutes one of the strongest controlled, non-focal perturbations of the human brain, offering a unique opportunity to study large-scale reorganization in vivo.

Neuroimaging suggests that ECT induces structural and functional changes in brain regions that are embedded in neural networks that have a causal role in treatment response^6^. These effects are often interpreted in the context of neuroplasticity, with proposed mechanisms including increased neurotrophin signaling^7^, neurogenesis^8^, synaptogenesis^9^, and glial proliferation^10^. These processes may arise from cytokine-induced transcriptomic and proteomic modifications^7,11^ and modulate neural signaling by affecting neurotransmitter and neurosteroid systems^12,13^. However, these components may not be universally necessary for treatment response, and evidence indicates that they may operate through partially independent pathways^9,14^. Despite these proposed mechanisms, the neurobiological and behavioral bases of ECT remain incompletely understood, particularly regarding how multilevel biological alterations translate into changes in human adaptive function and psychopathology. Previous meta-analyses and systematic reviews have examined structural and functional brain changes, as well as molecular correlates, but these efforts were limited by small samples, methodological heterogeneity, and a focus on single modalities^15-17^. A recent integrative review synthesized imaging, molecular, and mechanistic findings^18^, yet these insights remain fragmented. This lack of large-scale, multimodal, integrative studies makes it difficult to assemble these diverse findings into a coherent framework linking ECT-induced neurobiological effects with behavioral phenomena, symptom expression, and adaptive responses. In particular, how such a strong perturbation interacts with intrinsic network architecture, receptor distributions and cellular context remains largely unknown. Understanding these interactions is essential for identifying general principles by which large-scale perturbations reshape human brain organization.

Beyond mechanistic and behavioral insights, our work introduces a computational framework for integrating multiscale neurobiological data, enabling systematic biomarker discovery and benchmarking of neuromodulatory interventions while characterizing multiscale principles of perturbation-driven reorganization. Here, we present a multimodal investigation of the neurobiological substrates of ECT by leveraging large-scale neuroimaging data and applying enrichment analyses anchored in neurobiological ontologies. First, we identify reproducible hubs of ECT-related change across hundreds of treated subjects and map their embedding within distributed brain systems. We then use virtual histology to infer cellular contributors to the neuroplastic effects by testing for cell-type-specific enrichment. Leveraging electrophysiological data, we show that ECT-induced adaptations in network architecture align with intrinsic cortical dynamics. We then benchmark their spatial profile against positron emission tomography (PET)-derived chemoarchitectural maps of 17 neurotransmitter receptors and transporters to define their molecular context. We next use data-driven functional decoding to link ECT-induced adaptations to psychopathology-relevant symptom domains. Finally, leveraging a separate dataset of fast-acting antidepressant interventions, we use regularized multivariate modeling to quantify how ECT-induced adaptations relate to clinical improvement and to assess their treatment specificity. By integrating neuroplasticity, network dynamics, electrophysiology and neurotransmission with clinical change, this multiscale signature can inform individualized targeting and biomarker-guided treatment optimization in refractory psychiatric illness. More broadly, this framework provides a basis for identifying how intrinsic dynamics, receptor architecture and cellular composition constrain large-scale reorganization under strong neural perturbation.

## RESULTS

### ECT-induced morphological changes

We identified ECT effects on brain structure by meta-analyzing longitudinal voxel-based morphometry (VBM) data from ≈1,100 subjects across 45 experiments (Supplementary Fig. 1, Supplementary Table 1, Fig. 1a). Using activation likelihood estimation (ALE; Fig. 1b)^19,20^, we generated brain maps of ECT-induced morphological changes. ALE revealed convergence in an expansive right-hemispheric cluster covering temporal pole, temporoinsular transition zone, and amygdalohippocampal complex (Fig. 1c, Supplementary Table 3). Local maxima of convergence were also identified in the left amygdalohippocampal complex and in the ventral striatum (bilaterally) and subgenual cingulate cortex, two regions affected by depression and targeted by deep brain stimulation for treatment-resistant depression^21^. Notably, these effects were driven exclusively by volume increases.

**Fig. 1.**
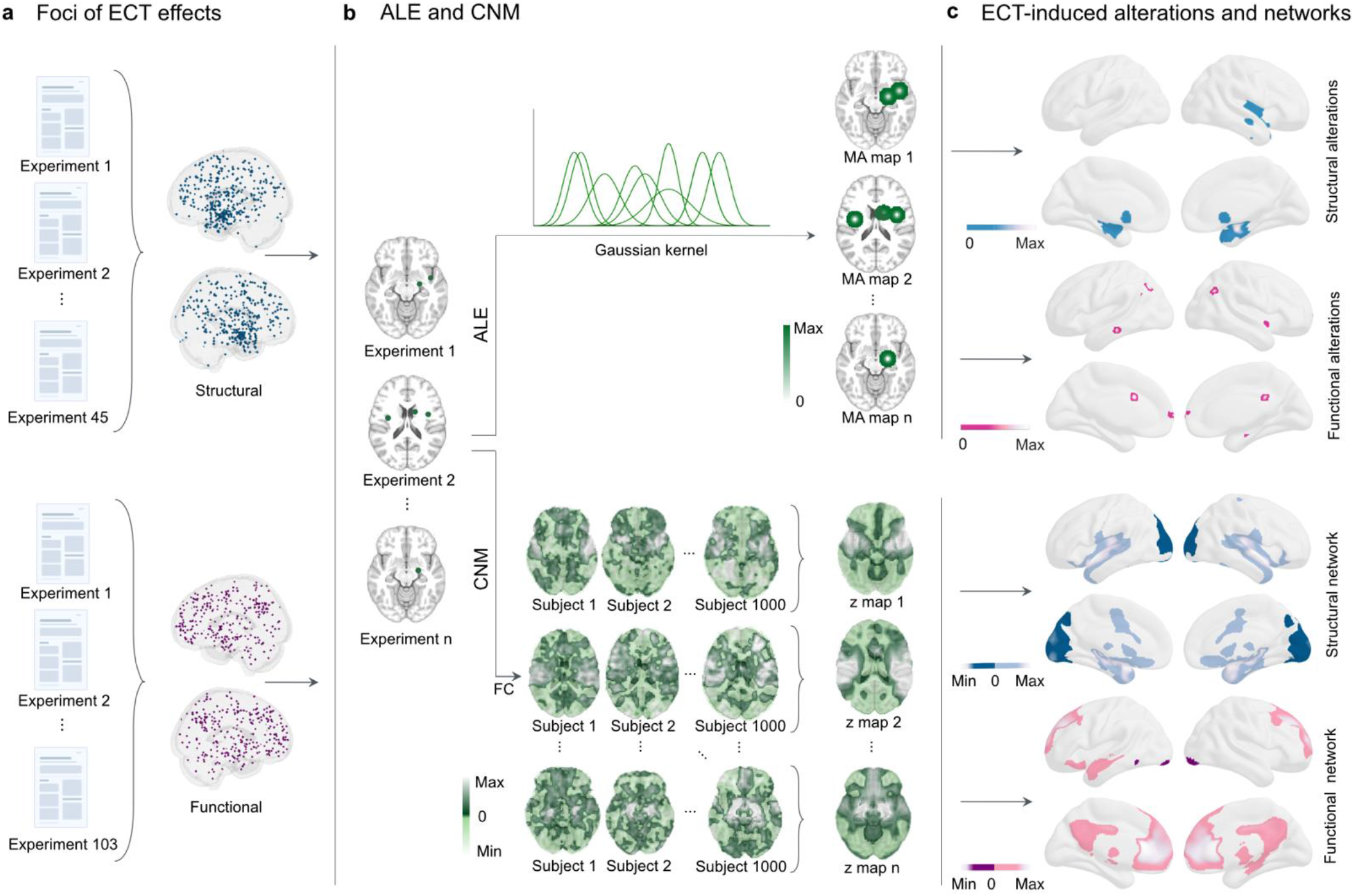
ECT-induced changes and networks. **a**, Pool of 408 foci (blue) representing ECT-induced structural alterations and 340 foci (purple) indicating functional changes, displayed on a transparent 3D brain. **b**, Workflow for activation likelihood estimation (ALE)^19,20^ and coordinate-based network mapping (CNM)^27^. Separate analyses were performed for structural and functional foci. Left: foci per experiment, where n indicates the number of experiments. Top: ALE models each experiment’s reported foci using Gaussian kernels scaled by sample size, generating modeled activation (MA) maps. Bottom: CNM computes, for each experiment, functional connectivity (FC) between foci and all other voxels using a normative connectome (N = 1,000), yielding n experiment-level mean Fisher *z*-maps. **c**, Top: ALE maps quantify convergence of changes across experiments at each location in the brain; significance assessed via threshold-free cluster enhancement (TFCE; 10,000 permutations). Three clusters of structural and six clusters of functional convergence (*p*<0.05, TFCE-corrected) are shown as surface projections. Bottom: CNM-derived networks were tested against zero using voxel-wise one-sample *t*-tests (TFCE; 10,000 permutations), identifying significant positive and negative brain networks of ECT-induced structural and functional changes (*p*<0.01, TFCE-corrected). These networks are visualized on the brain surface by connectivity direction, resulting in four distinct network maps.

How is this ECT-induced structural neuroplasticity mediated on a more granular level? To investigate the cellular basis, we applied a virtual histology^22^ approach to identify cell types enriched in regions modulated by ECT. We accessed gene expression distributions from the Allen Human Brain Atlas (AHBA)^23^, filtered them for biological relevance and consistency, and categorized them into nine major brain cell types (Fig. 2a). We identified significant positive associations between ECT-induced structural changes and interregional gene expression profiles specific to astrocytes (*r* = 0.29, *p*_perm_ = 0.001) and microglia (*r* = 0.30, *p*_perm_ = 0.001; Fig. 2b). Brain regions exhibiting greater structural change showed higher expression levels of astrocyte- and microglia-specific genes, suggesting a key role for glial plasticity in ECT-induced remodeling – consistent with peripheral biomarker evidence of ECT-induced glial proliferation^10^. We next compared the structural change map with an independent microstructure map^24^ using Spearman’s rank correlation and spatial autocorrelation-preserving null models^25^. ECT-induced morphological changes were greatest in regions with lower intracortical myelin (*r*=-0.34, *p*_spin_<0.001), in line with increased potential for synaptic plasticity (Fig. 2c)^26^.

**Fig. 2.**
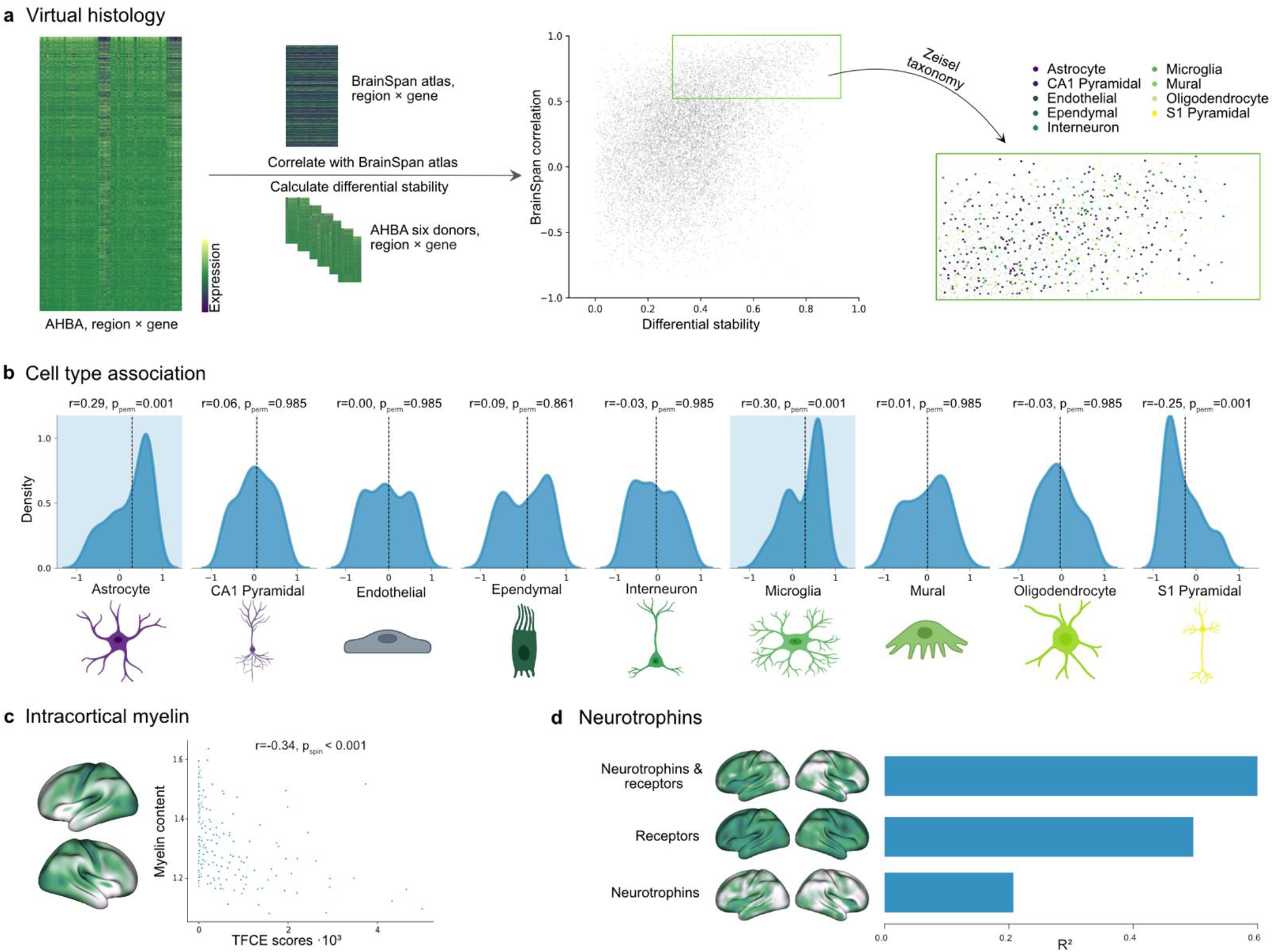
Molecular characterization of ECT-induced morphological changes. **a**, Workflow of the first part of the virtual histology^22^ approach to determine cell-type-specific gene distribution. Left: region-by-gene expression matrix including all gene distributions from the Allen Human Brain Atlas (AHBA)^23^ that passed intensity-based filtering. Middle: to ensure consistency across datasets, only genes with a correlation ≥ 0.52 (one-sided *p* ≤ 0.05) with an independent expression atlas (BrainSpan) were retained. To restrict the analysis to biologically relevant genes with consistent expression profiles across donors, the top 50% of genes with the highest differential stability were selected^31^. Right: genes passing both filtering steps were intersected with the Zeisel cell-type taxonomy^32^ and categorized into nine cell types. **b**, Spearman’s rank correlations between ECT-induced structural changes and expression profiles of cell-type-specific genes. The x-axis shows correlation coefficients; the y-axis represents estimated probability density. The vertical dashed line marks the average correlation coefficient for each cell type. Significant positive associations for astrocytes and microglia (p_perm_ ≤ 0.05, FWER-corrected) are highlighted with colored shading, indicating selective involvement of glial plasticity. **c**, Scatter plot showing the relationship between ECT-induced structural changes and intracortical myelin content^24^. Regions with lower myelin exhibited greater structural change (*r* = -0.34, *p*_spin_ < 0.001), consistent with increased potential for synaptic plasticity. The cortical distribution of myelin content is shown on the brain surface on the left. **d**, Ridge regression models predicting the spatial profile of gray matter change from gene expression distributions of (i) neurotrophins, (ii) their receptors, or (iii) both combined. Gene expression data were derived from the AHBA^23^. Model fits (*R*^*2*^) are shown in the bar plots and indicate stronger associations for receptor genes than for neurotrophins themselves, supporting the neurotrophic hypothesis. Brain surfaces show mean volumetric distributions of each gene set.

Finally, we examined whether neurotrophins – a family of neuroplasticity-promoting proteins^28^ – and their receptors, or both together, might regulate ECT-induced structural alterations. Using L2-regularized Ridge regression to capture multivariate correspondence, we fitted three models to predict the spatial profile of gray matter change from the gene expression distributions of (i) neurotrophins (*BDNF, NGF, NTF3, NTF4*), (ii) their receptors (*NTRK1, NTRK2, NTRK3, NGFR*), or (iii) both combined. The combined model showed the highest fit (*R*^2^ = 0.61), with receptor genes showing a stronger association (*R*^2^ = 0.49) than the neurotrophins themselves (*R*^2^ = 0.23), consistent with the neurotrophic hypothesis (Fig. 2d)^29^.

### ECT-induced changes in physiology and networks

To investigate the broader physiological and network-level effects of ECT, we extended our analysis to functional brain dynamics. We conducted ALE meta-analysis of longitudinal neuroimaging data examining ECT-induced functional brain changes from ≈1,200 subjects across 103 experiments (Supplementary Fig. 2, Supplementary Table 2, Fig. 1a). Convergent effects emerged in the left middle temporal and right insular cortices, bilaterally in the temporoparietal junction, the medial frontal pole, and the right hippocampus (Fig. 1c, Supplementary Table 4) – suggesting that the hippocampus may serve as a modulatory hub for structural and functional plasticity. We also observed a local maximum of convergence in the anterior midcingulate cortex, a known target in cingulotomy for treatment-resistant depression^30^. To determine whether ECT-related widespread brain changes across experiments localize to a common network, we applied coordinate-based network mapping (CNM)^27^ using a normative resting-state connectome (n = 1,000)^33^. CNM treats brain foci of ECT-induced changes as seed regions and computes their whole-brain functional connectivity, generating experiment-level network maps for longitudinal structural or functional changes. We identified regions showing positive or negative functional connectivity to these seeds across experiments, yielding common structural and functional networks (Fig. 1c). These results support the notion that ECT modulates large-scale brain dynamics that are disrupted in depression^34^ and schizophrenia^35^.

We next examined how these ECT networks relate to established functional brain systems using network correspondence analysis^36^. Spatial overlap was quantified with the Dice coefficient and assessed using spin-test permutation. To summarize correspondence across network definitions, we report the ten reference atlases with the highest (significant) Dice overlap for each ECT network map (Fig. 3a; full results in Supplementary Fig. 4). The positive structural network overlapped primarily with emotion/interoceptive, auditory, somatomotor, and divergent-cognitive systems. In contrast, the positive functional network corresponded most strongly to the default mode network and additionally engaged salience and parietal memory networks – systems critically implicated in psychiatric disorders^37-40^. Consistent with multimodal evidence of visual cortex disruptions – including altered connectivity in visual networks in patients with depression^41^ – the negative networks were linked to visual systems, while the structural network also engaged initiation and multiple-demand networks.

**Fig. 3.**
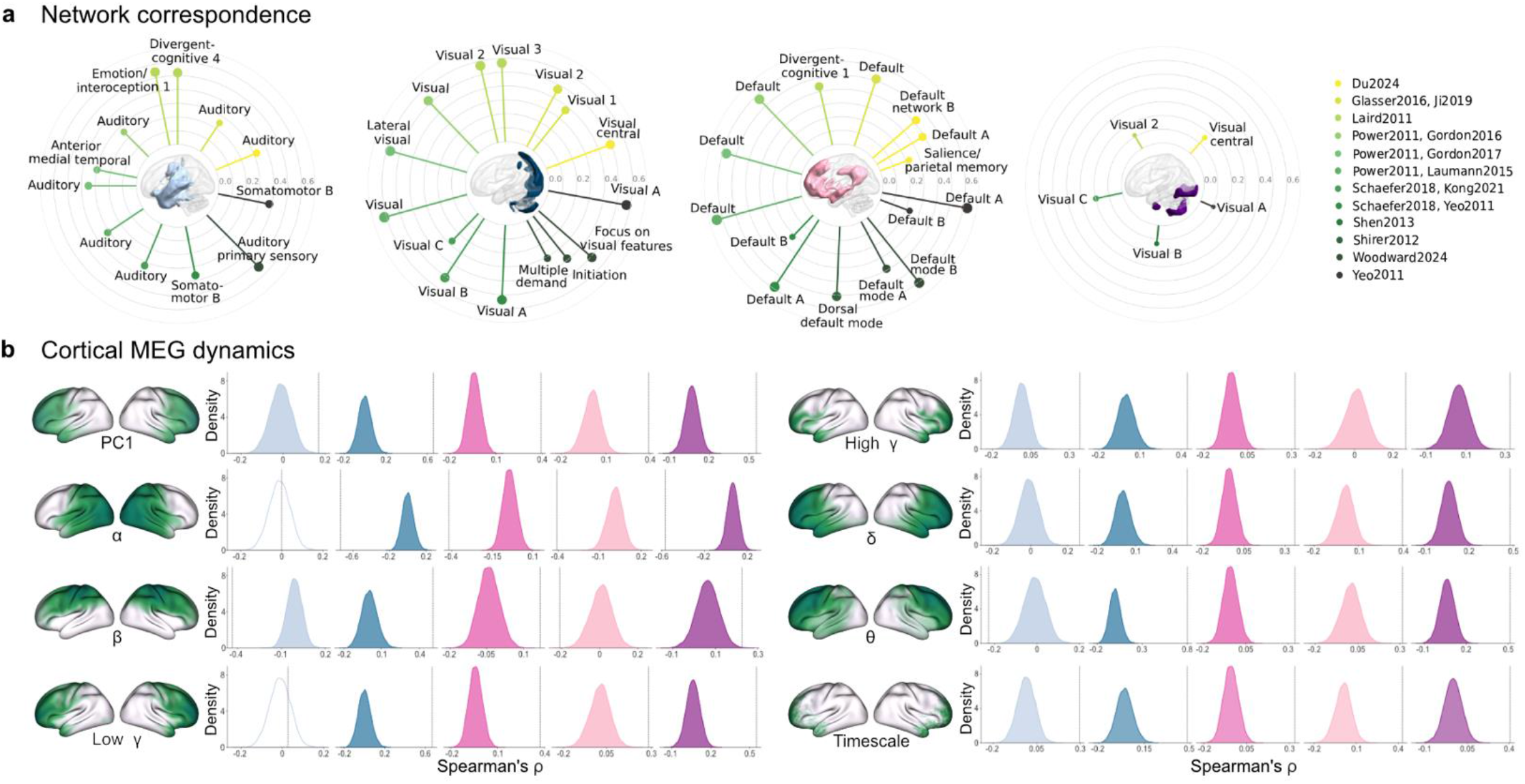
Network and electrophysiological correlates of ECT. **a**, Each ECT network map was analyzed using the Network Correspondence Toolbox^36^. From left to right, 3D brain renderings show the positive structural network (light blue), negative structural network (dark blue), positive functional network (light pink), and negative functional network (purple). Radial plots display Dice coefficients quantifying spatial overlap between each ECT map and canonical networks across multiple reference atlases (including ROI-, ICA-, and task-derived parcellations; see Supplementary Fig. 4 for full list and significance testing). For each map, results from the ten atlases with the highest Dice coefficients are shown, highlighting the most reproducible and spatially consistent overlaps. **b**, Spatial correlations between five ECT maps (four ECT network maps plus the functional change map) and eight electrophysiological metrics derived from MEG: PC1 (principal component summarizing global dynamics), six canonical frequency bands (α, β, low γ, high γ, δ, θ), and intrinsic timescale. The dashed line indicates the observed correlation coefficient; the curves represent spin-test null distributions preserving spatial autocorrelation, and shaded areas denote significant associations (*p* < 0.05, FWER corrected across all tests). Cortical distributions of MEG measures (derived from the HCP dataset^42^) are shown to the left of each plot. Color scheme as in a; functional changes are shown in pink. Thresholded ECT network maps are shown in panel a (cf. Fig. 1c for surface projections).

**Fig. 4.**
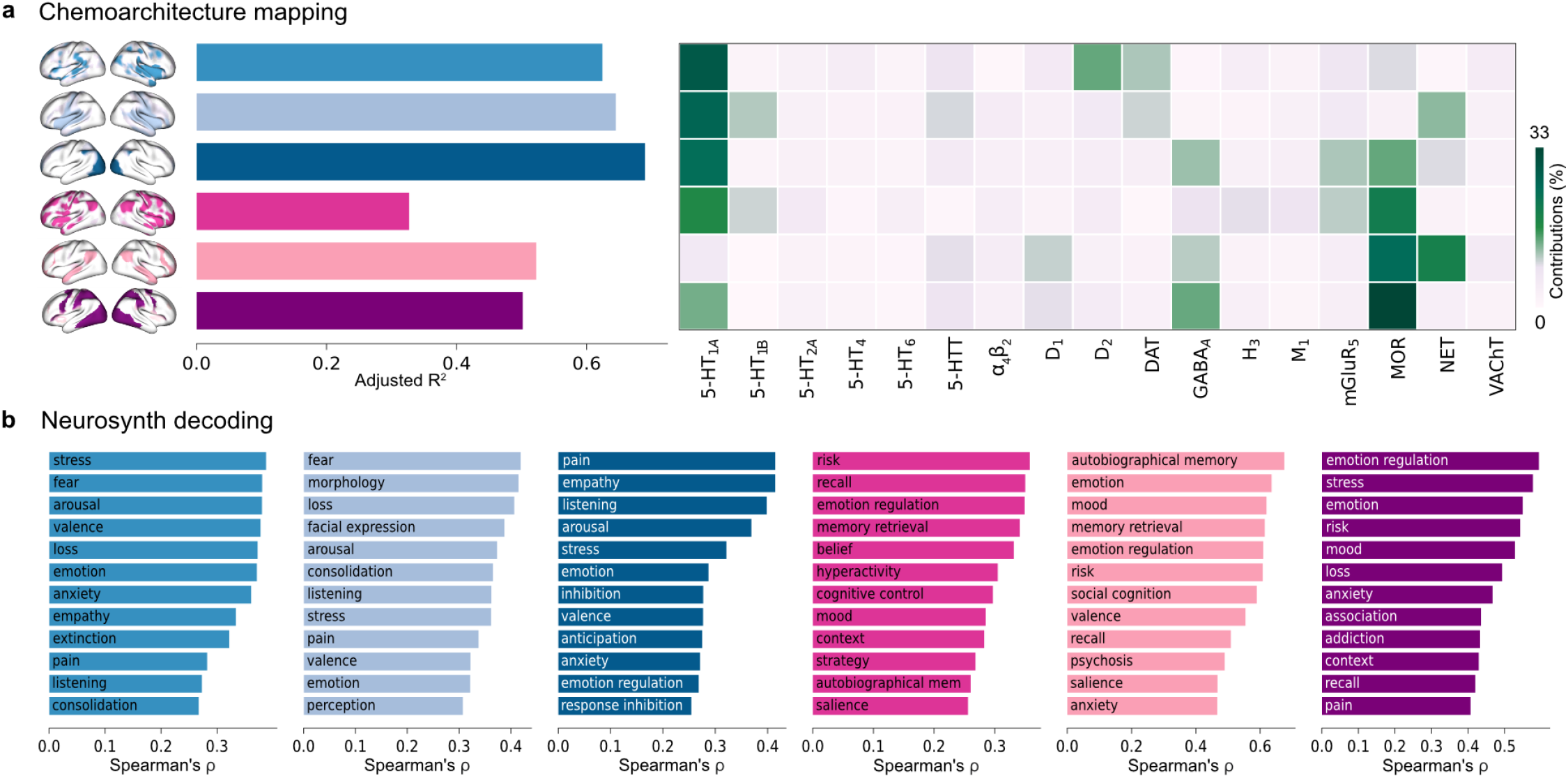
Chemoarchitectural and functional characterization of multimodal ECT effects. **a**, For each ECT map, a multiple linear regression model was fitted to predict its spatial profile based on 17 neurotransmitter receptor and transporter density maps across brain regions. ECT maps are shown on the left, with unthresholded versions from Fig. 1c displayed on the brain surface using the same color scheme: structural change (mid blue), positive structural network (light blue), negative structural network (dark blue), functional changes (pink), positive functional network (light pink), and negative functional network (purple). Model fits (adjusted *R*^2^) are presented in the bar plots. Dominance analysis was applied to estimate the relative contribution of each receptor and transporter to the model fit^46^; percent contributions are visualized in the heatmap. **b**, Each ECT map was correlated with 125 cognitive and behavioral meta-analytic activation maps from Neurosynth^47^. Bar plots show the top 10% of positive correlations, using the same color scheme as above. Full correlation profiles for all Neurosynth terms are provided in Supplementary Fig. 8. Abbreviations: mem, memory.

A comparison between all structural and all functional maps – using spatial autocorrelation-preserving null models and correction for multiple comparisons – revealed no significant spatial overlap, suggesting that ECT engages anatomically and functionally distinct mechanisms (Supplementary Fig. 5). Altogether, these findings highlight the spatially distinct modulatory effects of ECT on functional brain organization. To further characterize these functional effects, we examined whether ECT-related functional and network changes align with neural oscillatory rhythms derived from electrophysiological data – a more direct measure of intrinsic functional cortical dynamics. Specifically, we used magnetoencephalography (MEG) measures from the Human Connectome Project (HCP)^42^, including spectral power distributions across six canonical frequency bands and the intrinsic timescale, which reflects the temporal memory of a neural element. To identify a global pattern of cortical dynamics, we conducted a principal component analysis (PCA) across these seven electrophysiological measures, with the first principal component (PC1) explaining 69% of the total variance. We then correlated the ECT maps with PC1 and each individual electrophysiological measure, using spatial autocorrelation-preserving null models and correction for multiple comparisons. All ECT maps, except the positive structural network, showed significant alignment with PC1 and with all seven electrophysiological measures, indicating a shared global dynamic signature (Fig. 3b). Notably, the negative ECT brain networks exhibit particularly strong correlations with the global pattern of PC1 (*r* > 0.55) as well as with α, δ, and θ frequency bands (|*r*| ≥ 0.50), consistent with prior findings linking modulation of these rhythms to clinical improvement in psychiatric disorders^43-45^.

**Fig. 5.**
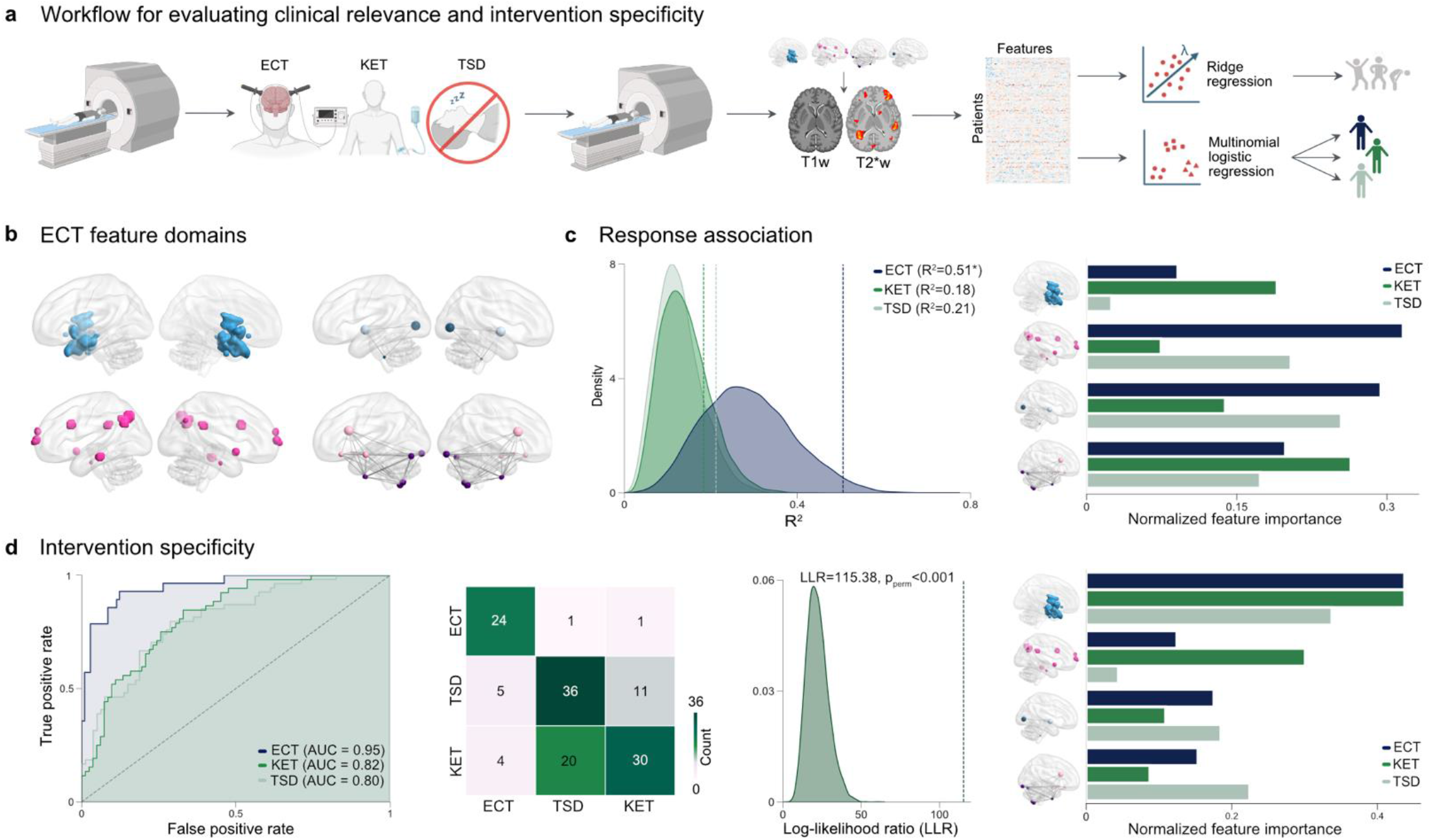
Assessing individual-level clinical relevance and intervention specificity across rapid-acting antidepressant interventions. **a**, A separate data set of fast-acting antidepressant interventions was analyzed, including depressed patients who underwent MRI and clinical assessment (Quick Inventory of Depressive Symptomatology; QIDS) before and after ECT, ketamine (KET) or total sleep deprivation (TSD) (n = 28/52/54). For each participant, pre-post differences in gray matter, fractional amplitude of low-frequency fluctuations (fALFF), and connectivity were extracted from ECT-derived feature domains, yielding a multimodal patient-by-feature matrix that was subsequently reduced using principal component analysis. Clinical relevance and treatment specificity were assessed using Ridge regression (top) and multinomial logistic regression (bottom). **b**, Four ECT feature domains were derived from thresholded ECT maps (Fig. 1c): structural regions of interest (ROIs; n = 5), functional ROIs (n = 11), connectivity among structural ROIs (n = 3 seeds) and among functional ROIs (n = 9 seeds). **c**, Ridge regression models were fit for each intervention group, using the first two principal components of each domain as predictors, with residualized change in QIDS as the target, controlling for age and sex. Left: permutation null distributions of *R*^2^ with observed values (dashed lines) for ECT (*p*_perm_ = 0.022), ketamine (*p*_perm_ = 0.209), and sleep deprivation (*p*_perm_ = 0.069). Asterisks denote significant effects, with strongest and only significant predictive performance observed for ECT, indicating an ECT-specific response signature. Right: normalized, magnitude-based (direction-agnostic) feature importance for each domain, computed in a balanced manner across predictors. Feature importance for demographic covariates is shown in Supplementary Fig. 9b; full regression coefficients appear in Supplementary Table 6. **d**, Treatment specificity. Multinomial logistic regression was used to classify intervention group from multimodal feature changes. Receiver operating characteristic (ROC) curves and one-vs-all area under the curve (AUC) demonstrate robust separability of ECT from other interventions, whereas ketamine and sleep deprivation show limited separability, as reflected in the confusion matrix. Permutation-based log-likelihood ratio (LLR) testing confirmed overall group separability (distribution under permutation with observed LLR values shown as a dashed line). Right: normalized, magnitude-based (direction-agnostic) feature importance for each domain. Feature importance for demographic covariates is shown in Supplementary Fig. 9c; full log-odds coefficients are provided in Supplementary Table 6.

### Mapping chemoarchitecture to multimodal ECT effects

Given the multimodal effects of ECT on the brain and the central role of neurotransmitter systems in modulating neuroplasticity and psychiatric symptoms, we examined the extent and specificity of their contributions to ECT-related brain changes. We used PET-derived distributions of 17 neurotransmitter receptors and transporters from recent PET atlases covering nine neurotransmitter systems in the human brain (Supplementary Table 5). For each ECT map, we fitted a multiple linear regression model to predict its profile based on receptor and transporter densities (Fig. 4a, left). Structural ECT maps yield the highest overall model fits (adjusted *R*^2^), indicating stronger alignment with underlying chemoarchitectural features than the functional maps.

We next applied dominance analysis to estimate the relative contribution (‘dominance’) of each receptor and transporter to the model fit (*R*^2^; Fig. 4a, right). The serotonin-1A receptor (5-HT_1A_) emerges as the dominant receptor across all ECT maps, with particularly strong predictive contributions in structural maps, including a peak in the map of structural changes. The 5-HT_1A_ receptor also exhibits high interactional dominance – defined as the increase in *R*^*2*^ when adding an independent variable to a model already containing all other predictors – suggesting it contributes unique, non-redundant information to the model predicting multimodal effects of ECT (Supplementary Fig. 7). These findings point to a prominent role of the 5-HT_1A_ receptor in the mechanism of ECT, consistent with PET evidence of ECT-induced alterations in postsynaptic 5-HT_1A_ binding potential^13^.

The dopamine D_2_ receptor ranked as the second most dominant predictor, with high interactional dominance in the map of structural changes, supporting prior PET findings of altered D_2_ binding following ECT^48^. In the functional maps, the µ-opioid receptor (µ) emerges as the strongest contributor to prediction of all ECT profiles, in line with evidence from animal models^49^. The norepinephrine transporter (NET) shows substantial contributions and high interactional dominance in both structural and positive functional ECT networks, while the GABA_A_ receptor contributes to both functional and negative structural networks. These findings suggest a network-mediated association between ECT effects and these neurotransmitter systems, which are implicated in the pathophysiology of depression^50,51^.

Collectively, these findings highlight the involvement of multiple neurotransmitter systems in modulating ECT effects on the brain, with primary contributions from metabotropic receptors and transporters. Among these, the 5-HT_1A_ consistently emerges as the most dominant predictor across modalities.

### Mapping multimodal ECT effects to psychopathology

Next, we investigated which mental and symptom-relevant processes are associated with multimodal ECT effects. To functionally annotate each ECT map, we quantified its spatial correspondence with 125 functional meta-analytic activation patterns from Neurosynth^47^. Briefly, Neurosynth aggregates data from thousands of functional magnetic resonance imaging (fMRI) studies to identify regional activations associated with specific cognitive functions and processes. We report the top-ranked positively associated terms (top decile, 12 terms) in Fig. 4b and provide the full set of correspondence statistics in Supplementary Fig. 8. Across maps, the correspondence profile was dominated by affective and regulatory processes (e.g., ‘emotion’, ‘stress’, ‘fear/anxiety’, ‘mood’, ‘arousal’, ‘empathy’, ‘emotion regulation’), suggesting that ECT exerts a multifaceted impact on affective brain systems. The functional ECT maps also align with memory-related terms such as ‘autobiographical memory’, ‘memory retrieval’ and ‘recall’, domains known to be affected by ECT^52^. Additional high-ranking annotations mapped to valuation and salience-related domains (e.g., ‘risk’, ‘valence’, ‘loss’, ‘salience’), somatosensory/interoceptive domains (‘pain’), and – within the same top-ranked set – psychosis-relevant symptom content (‘psychosis’). Together, these annotation position multimodal ECT effects at the intersection of affective regulation, valuation/salience, memory, and interoceptive symptom dimensions – including psychosis-relevant content.

### Response association and ECT specificity

Finally, we asked whether multimodal ECT-induced brain adaptations are clinically relevant and intervention-specific. Using a separate dataset of fast-acting antidepressant interventions (ECT, ketamine, and total sleep deprivation), we extracted four ECT-derived feature domains from the thresholded maps (Fig. 1c): gray-matter volume in structural ROIs, fractional amplitude of low-frequency fluctuations (fALFF) in functional ROIs, and connectivity within structural and functional networks (Fig. 5a). Positive and negative subnetworks were merged into integrated connectivity profiles. Treatment-induced changes were computed as pre-post differences and summarized via domain-wise principal component analysis (first two components per domain based on scree plot inspection; Supplementary Fig. 9a), to balance their contribution, reduce dimensionality and mitigate overfitting.

To quantify multivariate correspondence between feature-domain changes and clinical outcome (residualized change in Quick Inventory of Depressive Symptomatology (QIDS) scores), we used L2-regularized Ridge regression with permutation-based inference, fit separately within each intervention group. This regularized statistical-learning approach revealed the strongest – and only significant – within-sample association for ECT (*R*^2^ = 0.51, *p*_perm_ = 0.022), indicating an ECT-specific response signature (Fig. 5b). Feature importance patterns differed across interventions, suggesting distinct neural substrates of symptom improvement. In the ECT group, functional and connectivity changes contributed most to the model, consistent with a prominent role of network-level adaptations in rapid clinical response.

To assess intervention specificity, we tested whether patterns of multimodal change were separable across treatment groups, i.e., reflect treatment-specific biological effects rather than general markers of improvement, using multinomial logistic regression. Multimodal change profiles showed clear separability of ECT from the other interventions, with gray-matter alterations contributing most to group separability. Permutation-based log-likelihood ratio testing confirmed a significant group effect (Fig. 5c), indicating that ECT operates via a distinct mechanism of action in these domains. Together, these results indicate that the multimodal ECT signature is closely linked to symptomatic improvement and exhibits intervention-specific multivariate structure.

## DISCUSSION

Linking treatment-induced brain changes to clinical outcomes remains a key challenge in neuropsychiatry. In this study, we leveraged large-scale multimodal neuroimaging data from ≈2,300 subjects to delineate the neurobiological mechanisms underlying ECT-induced brain adaptations and their role in what confers clinical improvement. We generated spatial maps of ECT-induced brain changes, revealing hubs of structural, functional, and network-level reorganization. These adaptations appear to span multiple domains – including glial plasticity, neurotrophin signaling, and neurotransmitter systems – align with intrinsic oscillatory dynamics and emotion-related circuits, and predict clinical response, reflecting treatment-specific neural changes. Because ECT represents one of the strongest controlled perturbations of the human brain, these signatures may also reflect broader organizational constraints governing large-scale neurobiological reconfiguration. These findings demonstrate that large-scale perturbation reshapes multilevel brain organization in ways that are tightly linked to behaviorally relevant processes.

Decades of research have proposed diverse mechanistic hypotheses for ECT, implicating a range of neurobiological and physiological processes. By integrating large-scale multimodal data across psychiatric disorders, our work unifies these previously fragmented mechanistic theories into a data-driven, systems-level framework, offering a translational blueprint for understanding and optimizing therapeutic brain interventions.

We observed consistent volume increases in subcortical regions, including the hippocampus, amygdala, subgenual cingulate cortex, ventral striatum, and temporoinsular transition zone. Hippocampal enlargement represents the most consistent structural effect of ECT, with meta-analytic evidence also supporting amygdala changes^17,18,53^. These limbic regions – typically reduced in volume and functionally impaired in depression, schizophrenia, or bipolar disorder^54-61^ – are embedded in emotion/interoceptive and divergent-cognitive networks, converging on affective and cognitive systems^62^. In contrast, negative connectivity with visual networks highlights circuits disrupted in depression^41^. Such distributed engagement underscores how large-scale structural plasticity intersects with circuits central to adaptive emotional regulation and symptom expression. Additional links to sensorimotor networks suggest a distributed pattern of functional reorganization beyond primarily affective circuits. Taken together, these patterns indicate that the multiscale effects of ECT are embedded within large-scale systems supporting core psychological functions, including affective regulation, interoception, and memory – domains prominently disrupted across psychiatric disorders.

Collectively, these changes may reflect normalization of pathological circuits governing emotion-related processes, as indicated by Neurosynth analyses and co-localization with 5-HT_1A_ receptor density – a key modulator of emotional processing and pharmacological target in depression^63^. Structural alterations associated with the D_2_ receptor, central to schizophrenia pathophysiology, may signal improvement in positive symptoms^64^. The clinical relevance of these networks is underscored by neuromodulatory interventions such as deep brain stimulation of the subgenual cingulate cortex and ventral striatum^21^, and non-invasive transcranial magnetic stimulation targeting subgenual circuits^65^. Nevertheless, the link between structural changes and clinical response remains inconsistent across studies^18^. Such variability underscores the importance of considering how intrinsic dynamics, receptor gradients and cellular composition jointly constrain the expression of perturbation-induced effects.

We next examined the cellular and molecular processes underlying ECT-induced structural changes. Regions showing the strongest effects exhibited higher expression of astrocyte- and microglia-specific genes, suggesting that ECT promotes glial proliferation – consistent with peripheral biomarker evidence^10^ and preclinical findings^10,66^. Indirect indications of enhanced synaptic plasticity were reflected in reduced intracortical myelin^26^ and supported by animal models^67^, although one small pilot study reported divergent results^68^. Experimental work involving animal models on electroconvulsive shock (ECS) further demonstrates extended neuronal survival, increased dendritic complexity, and higher spine density^69^, indicating robust synaptogenesis^70^. ECS also induces angiogenesis and endothelial proliferation^71^, as well as neurogenesis in the dentate gyrus^72,73^ and frontal brain regions^74^ following repeated stimulation. While limited in vivo^75^ and post-mortem evidence^76^ supports ECT-induced neurogenesis, converging data from animal models^9,77^ and the low rate of adult neurogenesis^78^ suggest that generalized neural plasticity is the primary mechanism.

Additional evidence for neuroplasticity comes from ECT-related enhancement of neurotrophin signaling, with upregulation observed in animal models^79^ and a trend toward increased BDNF levels in humans^18^, while NT-3^80^ and NGF^80,81^ remain unchanged in small samples. Although disrupted neurotrophin pathways are implicated in depression^82^, schizophrenia^83^, and bipolar disorder^84^, the extent to which ECT-mediated modulation translates into clinical benefit remains uncertain^18^. Rather than simple changes in neurotrophin levels, our findings point to receptor-level adaptations as key mediators of neurotrophin signaling and downstream plasticity, paralleling classical neurotransmitter models^85^. Animal studies reported ECS-induced increases in phosphorylated TrkB and downregulation of full-length TrkB receptors^86^, with limited evidence linking BDNF-TrkB upregulation to symptom improvement^87^. However, data on ECT’s impact on neurotrophic receptors and downstream signaling remain sparse, underscoring the need for mechanistic studies.

Taken together, these neurotrophin-related changes and microglial increases^88^ – along with evidence of ECT-induced microglial activation^89,90^ – support the hypothesis that potentiated inflammatory processes contribute to BDNF release and, in turn, structural plasticity^7^. This neuroplasticity-neuroinflammation link is further corroborated by reports of ECT-induced upregulation of pro-inflammatory cytokines^7^ and metabolic shifts, including increased *N*-acetylaspartate and reduced Glutamate+Glutamine concentrations^91^. Similarly, astrocytic activation may amplify neuroplasticity-promoting pathways, consistent with their role in BDNF-TrkB signaling^10,89,92^. A recent preclinical study links these neuroinflammatory and neuroplastic mechanisms to ECT’s rapid antidepressant effects by implicating adenosine signaling as a pivotal mediator^93^.

Because psychiatric symptoms cannot be explained by structural alterations alone, and ECT likely exerts functional effects, we next examined its impact on functional reorganization. The hippocampus emerged as a central hub, showing structural and functional changes consistent with its multimodal involvement across psychiatric disorders^54,57-59^. Additional changes were observed in regions implicated in schizophrenia or bipolar disorder^94,95^, including the insular cortices and the temporoparietal junction. ECT-induced changes in the medial frontal pole and insula, together with positive connectivity to the default mode and salience networks, suggest an alignment of structural and functional reorganization, with these hubs integrating limbic input^96-98^. Dysregulation of these circuits is a hallmark of depression^38,98,99^ and schizophrenia^39^; our findings are consistent with rebalancing of their functional balance, supported by direct resting-state functional magnetic resonance imaging evidence^100^ and consistent with NET transporter modulation observed in animal models^101^ and clinical studies^102^.

Functional ECT maps aligned not only with emotion-related processes but also with memory-related terms such as ‘autobiographical memory’, ‘memory retrieval’, and ‘recall’. These associations were strongest for positive networks, reflecting their overlap with the parietal memory network and the default mode network – key system for autobiographical memory^103^. Memory impairment remains the most frequent cognitive side effect of ECT, typically involving anterograde and retrograde amnesia and deficits in autobiographical memory^52,104^. However, this alignment cannot be unequivocally attributed to adverse effects or therapeutic mechanisms, as ECT may also engage memory-related networks^100,103^ and improve cognitive function in depression. ECT maps further showed strong associations with µ-opioid receptor expression. Small studies using naloxone, a µ-opioid receptor antagonist, report no effect on retrograde memory impairment following ECT^105,106^, supporting a role for µ-opioid receptors in therapeutic network reorganization rather than in cognitive side effects. This interpretation aligns with the receptor’s involvement in affective regulation and stress buffering^107^, and its dysregulation in depression^107^ and schizophrenia^108^.

Negative structural and functional networks aligned with visual systems and altered GABA_A_ concentrations, consistent with prior reports of disrupted visual networks^41^, abnormal GABA_A_ levels in visual regions, and impaired GABA_A_-visual processing associations^109^ in depression. Our findings also indicate coupling with gamma oscillatory dynamics, which are tightly linked to GABA_A_ergic inhibition in visual cortex^110^. Together, these results are consistent with modulation of visual GABA_A_-dependent mechanisms reflected in oscillatory activity. However, existing studies on occipital GABA_A_ changes following ECT are limited by small sample sizes, inconsistent findings, and lack of correlation with clinical outcome^111,112^.

Collectively, our results reveal multimodal ECT-induced brain adaptations across structural, functional, network, and molecular domains, with functional changes mirrored in widespread electrophysiological dynamics. These effects highlight local and network-level reorganization across the frequency spectrum. Importantly, potential therapeutic effects are consistently reflected across modalities in their association with emotion-related processes, converging on 5-HT_1A_ receptor density as a key molecular substrate. This distributed therapeutic signature parallels the complexity of brain and molecular abnormalities in psychiatric disorders, and is consistent with coordinated modulation across multiple levels and their interactions. When viewed broadly, these cross-scale correspondences may represent general principles of how large-scale perturbations reorganize human brain systems.

Beyond mechanistic insights, we evaluated whether multimodal ECT-induced brain signatures relate to clinical outcome and exhibit treatment specificity. Leveraging a separate dataset of fast-acting antidepressant interventions, we used a regularized statistical-learning framework to quantify multivariate correspondence between intervention-related brain changes and symptom improvement, and to test separability of change profiles across treatments. Multimodal alterations showed a robust association with symptom improvement only in ECT, defining a clinically relevant and treatment-specific neurobiological signature. Feature domain-importance analyses further indicated that rapid response is primarily driven by network-level and functional adaptations, whereas structural remodeling likely reflects slower, intervention-specific neuroplasticity. Together, these findings help reconcile inconsistencies^18^ in prior single-modality work and underscore the value of integrative, multimodal approaches for precision psychiatry. Nevertheless, the multiscale associations observed here are consistent with a general organizational framework in which intrinsic dynamics, receptor architecture and cellular composition jointly constrain the brain’s response to large-scale perturbation.

Overall, our findings provide a blueprint for future studies to integrate these modalities, their underlying mechanisms, and interactions within unified designs, enabling a more complete understanding of ECT’s mechanism of action. Combining receptor imaging with longitudinal structural and functional measures could clarify causal molecular-circuit sequences underlying clinical response. Convergence on specific molecular targets – 5-HT_1A_, µ-opioid, and GABA_A_ receptors – and evidence for BDNF-TrkB signaling offer entry points for mechanistic research and biomarker-guided strategies for treatment optimization or personalization. Because these insights emerge from ECT-specific, response-related adaptations – and given that ECT remains the most effective brain-stimulation therapy^113^ – this integrative framework provides a foundation for precision psychiatry, bridging mechanistic insights with predictive modeling to inform next-generation neuromodulatory and pharmacological interventions.

Several methodological considerations warrant attention. First, enrichment analyses are inherently associative and do not permit causal inference; interpretation should be supported by convergent evidence from experimental studies. Second, generalizability may be constrained by heterogeneity in patient populations, treatment protocols, and imaging modalities, underscoring the need for replication in diverse cohorts using harmonized designs.

In summary, our multimodal synthesis delineates a cross-scale signature of ECT that links structural, functional, network and molecular adaptations to intrinsic electrophysiology and symptom-relevant processes. By embedding perturbation-related changes within receptor gradients, cellular context and canonical networks, the work outlines organizational constraints that appear to shape how the human brain reconfigures under strong stimulation. These findings provide a principled basis for future experiments aimed at testing putative molecular-to-circuit sequences and their relevance for behavior. The same integrative pipeline can benchmark neuromodulatory and pharmacological interventions against a common multiscale reference, enabling biomarker-guided personalization and comparative mechanism mapping across disorders. More broadly, the regularities identified here may extend beyond ECT to other forms of large-scale perturbation, offering a compact set of organizational principles for human brain adaptation.

## METHODS

### Data selection

Our study was conducted in accordance with the “Preferred Reporting Items for Systematic Reviews and Meta-Analyses (PRISMA)” statement and best practice guidelines for neuroimaging meta-analysis^114,115^.

A principled procedure to identify the relevant experimental studies published until December 31, 2024, was employed (Supplementary Fig. 1 and 2). We selected studies through two independent standard searches in the PubMed (https://www.ncbi.nlm.nih.gov/pubmed/) and ISI Web of Science (https://www.webofknowledge.com) databases. Regarding structural imaging, we used the term “electroconvulsive therapy” in combination with “MRI”, “structural MRI”, “structural magnetic resonance”, “gray matter”, “grey matter”, “VBM”, “voxel-based morphometry”, “neuroimaging”, or “imaging”. Regarding functional imaging, we used the term “electroconvulsive therapy” in combination with “resting-state”, “brain activity”, “arterial spin labeling”, “ASL”, “regional homogeneity”, “ReHo”, “glucose metabolism”, “single photon emission computed tomography”, “SPECT”, “positron emission tomography”, “PET”, “regional cerebral blood flow”, “rCBF”, “amplitude of low-frequency fluctuations”, “ALFF”, “degree centrality”, “blood oxygen level dependent”, “BOLD “, “functional connectivity density”. Further studies were found by means of the “related articles” function of the PubMed database and by tracing the references from the identified papers and review articles. Neuroimaging experiments were considered relevant when they reported longitudinal comparisons of gray matter or brain function before vs. after ECT. Only experiments reporting results of whole-brain group analyses with coordinates referring to a standard reference space (Talairach-Tournoux or Montreal Neurological Institute (MNI)) were included. Results of region-of-interest or seed-based connectivity analyses and studies not reporting stereotaxic coordinates were excluded.

On the basis of these search criteria, 25 papers employing voxel-based morphometry and 40 papers employing functional brain imaging techniques were found to be eligible for inclusion into the meta-analyses (Supplementary Table 1 and 2). The experimental contrasts involved increases (‘before < after ECT’) and decreases (‘before > after ECT’) of gray matter or brain function from longitudinal comparisons as well as analyses on main time or interaction effects indicating ECT-induced changes in gray matter or brain function.

Differences in coordinate spaces (Talairach vs. MNI space) were accounted for by transforming coordinates reported in Talairach space into MNI coordinates using a linear transformation.

Convergence of reported foci was analyzed for the main effects of ECT-induced changes in gray matter (45 experimental contrasts, 408 foci; Fig. 1a,c) and brain function (103 experimental contrasts, 340 foci; Fig. 1a,c), respectively. The denoted sample sizes (i.e., number of experimental contrasts) have been shown to be sufficient to achieve robust meta-analytic estimates^116^.

### Activation likelihood estimation

All statistical analyses were carried out using the revised ALE algorithm for coordinate-based meta-analysis of neuroimaging results^19,20^. This algorithm aims to identify regions with a convergence of reported coordinates across experiments that is higher than expected from a random spatial distribution. Foci are treated as centers of 3D Gaussian probability distributions capturing the spatial uncertainty associated with each focus^19^. Here, the between-subject variance is weighted by the number of participants per study, since larger sample sizes should provide more reliable approximations of the “true” activation effect and should therefore be modeled by more “narrow” Gaussian distributions. Subsequently, probabilities of all foci reported of a given experiment were combined for each voxel, yielding a modeled activation (MA) map^20^. Notably, foci were organized by subject group, which prevents multiple foci from a single experiment from cumulatively influencing MA values^20^. This approach hence prevents multiple experiments performed by one subject group from cumulatively influencing ALE values. It can thus be ruled out that effects are amplified by nonorthogonal contrasts (i.e., from the same study) being submitted to the same analysis. The union across these individual MA maps was then calculated to obtain voxel-wise ALE scores, i.e., an ALE map, which quantified the convergence across experiments at each location in the brain. More precisely, ALE scores were calculated by taking the voxel-wise union of their probability values, i.e., as:

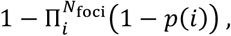

where *p*(*i*) is the probability associated with the *i*th focus at this particular voxel. To distinguish “true” from random convergence, ALE scores (i.e., the ALE map) were compared to an empirical null distribution of random spatial associations between experiments using a non-linear histogram integration algorithm^19^. The resulting random-effects inference focuses on the above chance convergence across studies rather than the clustering within a particular study^117^. This null hypothesis was derived by computing the distribution that would be obtained when sampling a voxel at random from each of the MA maps and taking the union of these values in the same manner as for the (spatially contingent) voxels in the original analysis^19^. The *p*-value of a “true” ALE score was then given by the proportion of equal or higher values obtained under the null distribution. The resulting nonparametric *p*-values were then assessed employing threshold-free cluster enhancement using 10,000 permutations for correcting multiple comparisons^19^.

Contributions were estimated by determining for each included experiment, how much it contributes to the summarized test-value (i.e., the ALE score) of a specific cluster. This was done by computing the ratio of the summarized test-values of all voxels of a specific cluster with and without the experiment in question, thus estimating how much the summarized test-value of this cluster would decrease when removing the experiment in question^114^.

For anatomical labeling, we capitalized on cytoarchitectonic maps of the human brain provided by the Statistical Parametric Mapping (SPM) Anatomy Toolbox (JuBrain Anatomy Toolbox v3.0)^118-120^. Clusters were thus assigned to the most probable histologically defined area at the respective location. This probabilistic histology-based anatomical labeling is reported in the Supplementary Tables 3 and 4.

References to details regarding cytoarchitecture are given in the table notes. For areas not listed in this tool, we relied on the Talairach Daemon^121^, including Structural Probability Maps, which in turn are based on 50 or more MRI brain volumes that were automatically labeled using the non-linear image-matching ANIMAL algorithm^122^.

### Gene expression data

Gene expression data were obtained from the Allen Human Brain Atlas (AHBA, https://human.brain-map.org)^23^, comprising six adult donors (ages 24–57 years; five males), with samples from the left hemisphere in all donors and from both hemispheres in two donors (ages 24 and 39 years; both male). Data processing was conducted using the *abagen* toolbox^123^ (v0.1.3; https://github.com/rmarkello/abagen) in volumetric MNI space, employing an extended version of the Desikan-Killiany atlas^124^ that includes 33 cortical and additional 8 subcortical regions per hemisphere.

Microarray probes were reannotated using updated mappings^125^, and probes lacking valid Entrez IDs were excluded. For genes represented by multiple probes, we selected the probe with the highest differential stability^31^ – defined as the consistency of regional variation across donors – to ensure robust gene-level estimates. Differential stability was calculated as:

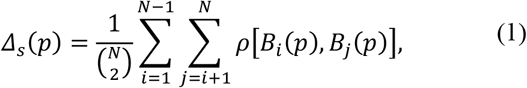

where *ρ* is Spearman’s rank correlation of the expression of a single probe *p* across regions in two donors *B*_*i*_ and *B*_*j*_, and *N* is the total number of donors. Regions correspond to anatomical designations defined in the AHBA ontology.

The MNI coordinates of tissue samples were updated using non-linear registration via Advanced Normalization Tools (ANTs; https://github.com/chrisfilo/alleninf). Samples were assigned to brain regions in the atlas if their MNI coordinates were within 2 mm of a given parcel. For regions without directly assigned tissue samples, each voxel was mapped to the nearest tissue sample from the donor to generate a dense, interpolated expression map. The average expression across all voxels in the region was then computed, weighted by the distance between each voxel and its assigned sample, to estimate parcellated expression values. Tissue samples not assigned to any brain atlas region were excluded from further analysis.

To address inter-subject variability, expression values for each tissue sample were normalized across genes using a robust sigmoid function^126^:

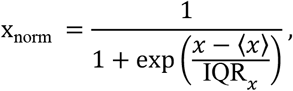

where ⟨*x*⟩ is the median and IQR_*x*_ is the normalized interquartile range of the expression of a single tissue sample across genes. Normalized expression values were then rescaled to the unit interval:

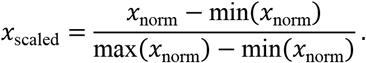

Gene expression values were subsequently normalized across tissue samples using the same robust sigmoid and rescaling procedure described above. For each donor, expression values from samples assigned to the same brain region were averaged. These regional values were then averaged across donors, resulting in a parcellated gene expression matrix with 41 rows per hemisphere (corresponding to brain regions) and 20,232 columns (corresponding to the retained genes).

### Virtual histology

To investigate how the meta-analytically derived spatial map of ECT-induced neuroplastic brain changes relates to cell-specific gene expression, we applied a virtual histology approach^22,127^. Gene expression data were processed as described above. Probes with intensity values below background noise in ≥ 50.00% of samples across donors were excluded^128^, resulting in 15,633 retained genes. Analyses were conducted on the left hemisphere (six donors) due to higher data quality, with the right hemisphere (two donors) used for validation.

A two-stage gene filtering process was applied to achieve consistency and representativeness of interregional gene expression profiles. First, we selected the top 50% of genes with the highest differential stability as defined in Equation (1), to retain only biologically relevant genes with consistent expression profiles across donors^31^. Second, to ensure cross-dataset consistency, we retained only genes with a Spearman correlation of ≥ 0.52 (one-sided *p* ≤ 0.05) with an independent gene expression atlas (BrainSpan, www.brainspan.org) across 11 overlapping regions were retained. This yielded 2,428 genes from an initial subset of 7,816.

These genes were then intersected with single-cell RNA sequencing data from the mouse hippocampus and primary somatosensory cortex (S1)^32^. Genes present in both datasets were categorized into nine canonical cell types: CA1 pyramidal (99 genes), S1 pyramidal (81 genes), interneurons (92 genes), astrocytes (59 genes), microglia (50 genes), oligodendrocytes (60 genes), mural cells (23 genes), endothelial cells (46 genes), and ependymal cells (73 genes). To assess the relationship between the spatial brain map and cell-type-specific gene expression, we computed Spearman’s rank correlations between the spatial map and the expression profile of each gene across regions of the extended Desikan-Killiany atlas. For each cell type, the mean correlation coefficient was calculated by averaging correlations across all genes assigned to that cell type. Statistical significance was assessed using a two-sided test against an empirical null distribution generated from 10,000 random gene sets (matched in size to each cell type) drawn from the 2,428-gene reference panel. Results for the left hemisphere are shown in Fig. 1b; right hemisphere validation is presented in Supplementary Fig. 3. All 18 tests were FWER-corrected for multiple comparisons^129^.

### Myelin

To estimate intracortical myelin content, we used T1-weighted/T2-weighted (T1w/T2w) signal intensity ratios from 417 unrelated participants (ages 22–37 years; 193 males) from the Human Connectome Project (HCP, S1200 release)^42,130^. Group-level intracortical myelin maps^24^ were derived using the *neuromaps* software package (https://github.com/netneurolab/neuromaps)^25^, which provides tools for accessing, transforming, and parcellating curated brain maps from the literature. The T1w/T2w (myelin) map was generated in the fsLR32k surface space and subsequently parcellated according to the Craddock (CC200) atlas^131^.

### Spatial null models

To assess the statistical significance of spatial associations across brain regions while accounting for spatial autocorrelation, we employed spatial permutation tests (“spin tests”)^132,133^. For tests implemented via the *neuromaps* package^25^, a surface-based representation of the brain parcellation was created on the FreeSurfer *fsaverage* surface using files from the Connectome Mapper toolkit (https://github.com/LTS5/cmp). A spherical projection of the surface was used to define spatial coordinates for each parcel by identifying the vertex closest to the parcel’s center of mass^134^. Parcel coordinates were then randomly rotated on the sphere, and each original parcel was reassigned the value of the nearest rotated parcel. This procedure was repeated 10,000 times. To avoid upsampling, the spin test was performed at the parcel level rather than the vertex level, and separately for each hemisphere. Resulting *p*-values were FWER-corrected for multiple comparisons^129^.

### Network mapping

We employed a network mapping approach to determine whether heterogeneous peak foci across studies localize to a functionally connected, common network^27^. Spherical seeds with a 4 mm radius were created, centered at each reported focus^27^. To enable experiment-level random-effect inferences^47,117^, we merged the spheres from the same experiment and organized foci by subject group to obtain a combined seed. Using a publicly available normative connectome from 1,000 healthy individuals (GSP1000 Preprocessed Connectome for Lead DBS; https://dataverse.harvard.edu/dataverse/GSP), we identified brain-wide functional connectivity network patterns associated with each combined seed^33^. Specifically, for each subject in the GSP1000 dataset, we calculated the Pearson’s correlation coefficient between the average time course of all voxels within the combined seed and the time course of every voxel in the entire brain. The resulting subject-level correlation maps were transformed into Fisher *z-*maps and averaged across subjects to yield an experiment-level mean Fisher *z-*map.

To identify consistent connectivity patterns (i.e., networks connected to the combined seed) across experiments, we compared the mean Fisher *z*-maps against zero using voxel-wise one-sample *t*-tests in SPM12 software (SPM12, http://www.fil.ion.ucl.ac.uk/spm/software/spm12).

Nonparametric *p*-values were assessed using TFCE with 10,000 permutations, implemented via the SPM TFCE toolbox (https://github.com/ChristianGaser/tfce). Resulting maps were separated into positively and negatively connected networks, reflecting regions with positive or negative functional connectivity to the seeds.

### Network correspondence

To examine whether ECT-related networks align with established functional systems, we used the Network Correspondence Toolbox^36^. A broad set of reference atlases was included, covering ROI-based, ICA-derived, and task-based parcellations (e.g., Glasser2016, Schaefer, Yeo, Shen; full list in Supplementary Fig. 4). When multiple resolutions were available, finer-grained parcellations were prioritized to maximize anatomical specificity. The selected atlases included Glasser2016^24^ with Ji2019^135^, Laird2011^136^, Schaefer2018^137^ with Kong2021^138^, Schaefer2018 with Yeo2011^139^, Shen2013^140^, Shirer2012^141^, Power2011^142^ with Gordon2016^143^, Power2011 with Laumann2015^144^, Power2011 with Gordon2017^145^, Yeo2011, Woodward2024 (10.5281/zenodo.4274397), and Du2024^146^.

ECT-induced brain networks and atlas templates were projected into a common space (fsaverage6), and correspondence was assessed using the Dice coefficient (range: 0 = no spatial overlap, 1 = complete spatial overlap).

Statistical significance was assessed via spin permutation tests^132,147^ (1,000 iterations), which randomly rotate spherical projections of the ECT cortical network maps to generate null distributions while preserving spatial structure.

For each ECT network, we report spatial correspondence results from the 10 atlases with the highest Dice coefficients (Fig. 3a), highlighting the most reproducible and spatially consistent correspondence (full results in Supplementary Fig. 4).

### MEG data acquisition and pre-processing

Six-minute resting-state, eyes-open magnetoencephalography (MEG) recordings were acquired for 33 unrelated participants (ages 22–35 years; 17 males) from the HCP (S900 release)^42,130^. Full acquisition protocols are detailed in the HCP S900 Release Manual. For each participant, power spectra were computed at the vertex level across six different frequency bands: δ (2–4 Hz), θ (5–7 Hz), α (8–12 Hz), β (15–29 Hz), low γ (30–59 Hz), and high γ (60– 90 Hz), using the open-source software *Brainstorm*^148,149^.

Preprocessing included notch filtering at 60, 120, 180, 240, and 300 Hz, followed by a high-pass filter at 0.3 Hz to remove slow-wave and DC-offset artifacts. Preprocessed sensor-level data were projected to source space on the HCP’s fsLR4k cortical surface using overlapping spheres head models. Data and noise covariance matrices were estimated from resting-state and noise recordings, respectively. Source activity was reconstructed using Brainstorm’s linearly constrained minimum variance (LCMV) beamformer.

Power spectrum density (PSD) was estimated using Welch’s method with 4-second windows and 50% overlap. Average power within each frequency band was computed at each source vertex. Intrinsic timescales of MEG signals were estimated using spectral parameterization with the FOOOF (Fitting Oscillations and One Over F) toolbox^150,151^. Group-averaged source-level PSD maps and intrinsic time scale map, available on the fsLR4k surface, were downloaded from *neuromaps*^25^ and parcellated into the regions of the Craddock (CC200) atlas^131^.

### Neurotransmitter receptors and transporters

Neurotransmitter receptor and transporter density data were obtained using the *neuromaps* software package^25^. Volumetric positron emission tomography (PET)-derived density maps from healthy volunteers (Supplementary Table 5) were accessed and parcellated – along with the unthresholded source brain maps of ECT-induced effects – according to the Craddock (CC200) atlas^131^. All maps were *z*-scored prior to analysis. Following established methodology^152^, receptor and transporter types with multiple PET maps acquired using the same tracer (i.e., 5-HT_1B_, D_2_, mGluR_5_, and VAChT) were combined using a sample size-weighted average. Prior to averaging, we confirmed high inter-map correlations (Supplementary Fig. 6), thereby increasing the reliability of the resulting density estimates by leveraging larger sample sizes.

### Dominance analysis

To assess the relative contribution of each predictor (parcellated neurotransmitter distributions) to ECT-induced changes, we performed dominance analysis. As a preparatory step, all parcellated ECT maps were Box-Cox transformed^153^ with parameters *λ*_1_ = 0 and *λ*_2_ = 23. This transformation, commonly used in regression modeling, establishes a log-linear relationship between the predictors and the outcome. Given that dominance analysis employs various regression models, this transformation improves model validity by addressing regression assumptions and enhancing the reliability of metrics such as the coefficient of determination (*R*^2^).

Dominance analysis was conducted using the dominance-analysis Python package (https://github.com/dominance-analysis/dominance-analysis)^46^. This method quantifies the relative importance (“dominance”) of each independent variable by fitting all possible submodels of a multiple linear regression (i.e., 2^*p*^−1 models for *p* predictors). Total dominance is defined as the average of the relative increase in *R*^*2*^ when a given predictor is added to a submodel. The sum of the total dominance values across all predictors equals the *R*^*2*^ of the full model, allowing for normalization and interpretation as percentage contributions.

Interactional dominance refers to the increase in *R*^2^ when adding a predictor to a submodel that already includes all other predictors. This value is normalized by *R*^2^ of the full model and thus represents one term in the average that defines total dominance. A predictor with high interactional dominance contributes more uniquely in the presence of all other predictors, and vice versa for low interactional dominance, indirectly reflecting its shared variance with them.

Unlike approaches based solely on regression coefficients or univariate correlations, dominance analysis accounts for interactions among predictors and provides an interpretable partitioning of the overall model fit. This makes it particularly well-suited for comparing predictor importance both within and across models.

### Neurosynth

To infer the psychopathological relevance of ECT-induced brain changes, we used *Neurosynth*, a large-scale meta-analytic platform that aggregates functional brain-imaging results with rich experimental descriptions from over 14,000 published studies^47^. *Neurosynth* computes probabilistic association maps by identifying high-frequency terms (e.g., “pain”, “attention”) that are co-reported with standardized voxel coordinates of neural activity responses (https://github.com/neurosynth/neurosynth). These association maps estimate the probability that a given term is reported in a brain-imaging experiment, conditional on activation at a given voxel.

Because *Neurosynth* matches voxels to more than 1,000 terms – many of which are anatomical, disease-related, or otherwise non-psychological and difficult to interpret (e.g., “accumbens”, “adhd”, “arterial spin”) – we restricted our analyses to terms reflecting psychological processes. Following previous methodology^154^, we intersected the *Neurosynth* vocabulary with the *Cognitive Atlas* (https://www.cognitiveatlas.org/).^155^, a curated ontology of cognitive science. This yielded 125 terms spanning broad domains (e.g., “attention”, “emotion”), specific cognitive processes (e.g., “visual attention”, “episodic memory”), behaviors (e.g., “eating”, “sleep”), and emotional states (e.g., “fear”, “anxiety”). All terms are provided in Supplementary Fig. 8. The corresponding volumetric association test maps were parcellated according to the Craddock (CC200) atlas^131^ using the *neuromaps*^25^ package.

### Multimodal data for fast-acting antidepressant interventions

To investigate whether ECT-related brain changes observed in key regions of interest (ROIs) – derived from structural, functional, and network-level meta-analytic maps – are clinically relevant and treatment-specific, we analyzed multimodal MRI data from three cohorts of patients with major depression undergoing fast-acting antidepressant interventions: ECT, serial ketamine infusions, or total sleep deprivation (TSD).

A total of 153 patients participated:

- ECT (n = 33)
- Ketamine (n = 60)
- TSD (n = 60)

All participants underwent structural T1-weighted and resting-state T2*-weighted MRI scans at baseline and following treatment. Imaging protocols were harmonized with the Human Connectome Project for Aging (HCP-A), and all scans were acquired on the same 3T Siemens Prisma system at UCLA.

Patients met DSM-5 criteria for non-psychotic major depression and were experiencing a current episode of at least six months duration. Detailed inclusion and exclusion criteria, as well as clinical and behavioral assessments (e.g., QIDS^156^), are described in the PDC 1.0 reference manual (https://www.humanconnectome.org/storage/app/media/documentation/PDC1.0/PDC_1.0_Release_Manual.pdf).

Overall, 19 patients were excluded due to missing scans or clinical data, resulting in a final sample of 134 patients (ECT: n = 28, ages 20–75 years, 10 males; ketamine: n = 52, ages 20–59 years, 28 males; TSD: n = 54, ages 20–56 years, 24 males).

Structural T1-weighted images were processed using the standard longitudinal pipeline of the Computational Anatomy Toolbox (CAT12.9; https://neuro-jena.github.io/cat12/) implemented in SPM12, including intra-subject bias correction, tissue segmentation, and spatial normalization to MNI space. Longitudinal registration was performed to account for within-subject anatomical consistency across timepoints. Modulated normalized gray matter maps were smoothed with an 8 mm full-width at half-maximum (FWHM) Gaussian kernel. All images passed visual inspection and CAT12’s automated quality control metrics. For subsequent analyses, gray matter volume (GMV) was extracted for all predefined regions of interest (ROIs) using CAT12’s ROI-based morphometry framework.

Resting-state functional MRI data were preprocessed using the Data Processing Assistant for Resting-State fMRI (DPARSF^157^), following a standardized pipeline previously validated in large-scale neuroimaging studies (e.g., ref.^158^). Preprocessing steps included slice timing correction, realignment, normalization to MNI152 space, spatial smoothing with a 6 mm FWHM Gaussian kernel, nuisance regression (including Friston-24 motion parameters, white matter, and CSF signals), and temporal band-pass filtering (0.01–0.08 Hz).

Following preprocessing, we computed fractional amplitude of low-frequency fluctuations (fALFF) for each ROI, providing a voxel-wise measure of spontaneous neural activity. In addition, we constructed ROI-to-ROI functional connectivity matrices by calculating Pearson correlation coefficients between mean time series of ROIs within each ECT network map, yielding two modality-specific connectivity profiles for subsequent analysis.

For each modality, change scores were computed as pre-minus post-treatment values. Clinical outcome was quantified as residualized QIDS change to account for baseline.

### Principal component analysis

To identify global patterns of cortical dynamics, we first performed principal component analysis (PCA) across seven electrophysiological measures. PCA identifies orthogonal directions in feature space that capture maximal variance in the data, yielding a set of uncorrelated components^159^. We used the implementation in *scikit-learn* (https://scikit-learn.org) to perform the analysis.

The input data matrix *X* ∈ ℝ ^*n*×*p*^ contained seven electrophysiological features across *n* brain regions defined by the Craddock 200 (CC200) atlas^131^. Prior to decomposition, each feature was *z*-scored across brain regions to ensure comparability. PCA was performed via singular value decomposition (SVD), such that:

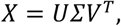

where *U* ∈ ℝ ^*n*×*n*^ and *V* ∈ ℝ ^*p*×*p*^ are orthonormal matrices, and *Σ* ∈ ℝ ^*n*×*p*^ is a diagonal matrix of singular values. The columns of *V* define the principal axes in feature space, and projecting *X* onto these axes yields the principal components: *XV*. The squared singular values (diagonal values of *Σ*^2^) are proportional to the variance explained by each component. The first principal component, which explained the largest portion of variance, was retained for further analysis.

Separately, we applied PCA to each ECT feature domain (GMV, fALFF, structural and functional network connectivity) to reduce dimensionality and capture dominant patterns of variation within each modality. For each domain, the input matrix *X* ∈ ℝ ^*n*×*p*^ comprised *p* features (ROIs or connectivity measures) across *n* patients. All features were *z*-scored across observations. PCA was performed via SVD as described above, and the first two principal components from each domain were retained for subsequent multivariate analyses.

### Ridge regression

Ridge regression was selected for its robustness in high-dimensional settings and its ability to handle multicollinearity while improving model stability. We applied Ridge regression in two analyses. First, we examined how neurotrophins and their receptors relate to ECT-induced structural brain changes. Prior to model fit, the parcellated meta-analytic ECT map was Box-Cox transformed using the same parameters as in the dominance analysis (*λ*_1_ = 0, *λ*_2_ = 23)^153^. Ridge regression was chosen to mitigate multicollinearity among predictors (variance inflation factors (VIF) ≤ 7.9) and overfitting by introducing an *ℓ*^2^ penalty to the ordinary least squares loss function^160^.

Second, Ridge regression was used to model multivariate associations between changes in the four ECT-derived feature-domains (plus age and sex) and residualized QIDS change, separately for each intervention group. We used ridge regularization to stabilize coefficient estimates, thereby reducing model variance and enabling more comparable domain-importance profiles across groups, particularly in the smaller ECT cohort. Model fit is reported as within-sample explained variance (*R*^2^) and assessed via permutation testing. Predictors were *z*-scored before fitting, and regression coefficients were estimated by solving the following numerical optimization problem:

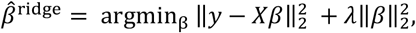

where *y* ∈ ℝ ^*n*^ is the outcome vector (ECT-induced structural changes across *n* brain regions, or clinical outcome across *n* patients), *X* ∈ ℝ ^*n*×*p*^ is the design matrix containing *p* predictors (gene-expression levels of neurotrophins and receptors, or ECT feature domains together with demographic covariates), and *β* ∈ ℝ ^*p*^ denotes the regression coefficients. The *ℓ*^2^-norm is defined as 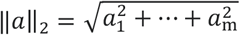. The regularization parameter *λ* ≥ 0 was optimized via cross-validation using the RidgeCV implementation from *scikit-learn* (100 logarithmically spaced values across [5 × 10^-3^, 5 × 10^9^]).

To assess feature importance of the ECT feature domains, we computed balanced contributions of the first two principal components per domain (PC1 and PC2) (*balanced importance*: 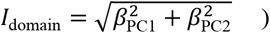, capturing the combined magnitude of both components while ensuring that domains remained influential even if one component dominated. All predictor coefficients, including balanced domain contributions, were converted to absolute values and normalized so that their sum equaled one within each model, facilitating comparability across intervention groups and preventing disproportionate weighting. Multicollinearity was minimal across all models (VIF ≤ 2), supporting interpretability of coefficient magnitudes and validity of this normalization approach.

Significance testing (10,000 iterations) was performed by randomly permuting clinical outcomes (residualized QIDS change) and refitting the Ridge model with the group-specific optimal regularization parameter, generating null distributions of *R*^2^ values.

### Multinomial logistic regression

To examine group-specific patterns in multimodal feature domains, we applied multinomial logistic regression to classify patients into one of three intervention groups by posing a three-class classification problem (ECT, ketamine, or total sleep deprivation). This approach generalizes binary logistic regression to multiclass problems by estimating a set of linear decision boundaries across all categories. Specifically, the probability that observation *i* belongs to class *k* is modeled as:

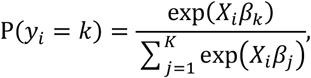

where *y*_*i*_ ∈{1, …,*K*} denotes the class label, *X*_*i*_ ∈ℝ^*p*^ is the predictor vector for subject *i*, and *β*_*k*_ ∈ ℝ ^*p*^ are the regression coefficients associated with treatment class *k*.

Model parameters were estimated by maximizing the log-likelihood function using the *limited-memory Broyden– Fletcher-Goldfarb-Shanno* (L-BFGS) optimization algorithm, as implemented in *scikit-learn*. To mitigate bias arising from class imbalance, inverse-frequency class weights were applied. All predictors were *z*-scored prior to analysis to ensure comparability of coefficients across features.

Estimated coefficients (*β*_*k*_) represent log-odds corresponding to predicting class *k*, with positive values increasing and negative values decreasing the odds of membership. Feature importance was computed in a direction-agnostic, domain-balanced, and normalized manner, consistent with the Ridge regression analysis.

To assess overall model significance, we performed a permutation-based likelihood ratio test. Specifically, we compared the log-likelihood of the fitted multinomial logistic regression model (*LL*_full_) to that of a null model containing only an intercept term (*LL*_null_). The log-likelihood was computed as:

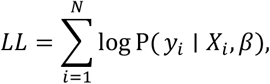

where P(*y*_*i*_ ∣ *X*_*i*_, *β*) is the predicted probability of observation *i* belonging to its true class. The observed likelihood-ratio statistic was calculated as *LR*_obs_ = −2 · (*LL*_null_ − *LL*_full_). To obtain an empirical *p*-value, we permuted class labels 10,000 times, refit the model for each permutation, and recomputed *LR*_obs_ to generate a null distribution.

## Data availability

Coordinates used in the meta-analyses and coordinate-based network mapping are available from the included studies referenced in the Supplement. Publicly available datasets include the Allen Human Brain Atlas (https://human.brain-map.org), BrainSpan (https://www.brainspan.org), GSP1000 Preprocessed Connectome (https://dataverse.harvard.edu/dataverse/GSP), and the Cognitive Atlas (https://www.cognitiveatlas.org/). MEG, neurotransmitter, and myelin maps are available through the *neuromaps* package (https://github.com/netneurolab/neuromaps/), and Neurosynth data are available through *Neurosynth* (https://neurosynth.org/). The PDC Release 1.0 data are accessible in the NIMH Data Archive (https://dx.doi.org/10.15154/00b0-hw43).

## Acknowledgements

None.

## Author contributions

S.K. and T.B.P. conceived the study. S.K. designed the analyses and implemented virtually all technical analyses; T.B.P. performed the meta-analyses and supervised the project. H.K.N. developed the initial pipeline for the network-correspondence analyses. M.B. preprocessed the original neuroimaging data. S.Z. collected data under the supervision of K.S. J.D. and J.Y.H. advised on neurotransmitter analyses; J.Y.H. additionally advised on Neurosynth analyses. B.M. advised on enrichment analyses. D.B. and S.B.E. advised on statistical-learning analyses; S.B.E. also provided software. T.C.B., C.N., M.G. and B.D. contributed to the study concept. S.K. prepared the first draft of the manuscript; T.B.P. provided the initial critical revision. All authors revised the manuscript and approved the final version.

## Competing interests

J.D. received support from the German Research Foundation (DFG; project number 549186835). D.B. is shareholder and scientific advisor of MindState Design Labs (USA) and Biossil (Canada); the contribution to this manuscript is unrelated to this affiliation. T.B.P. received support from the German Research Foundation (DFG; project number 519094028). Within the past three years, T.B.P. received honoraria for speaking engagements from Laboratorios Farmacéuticos Rovi; this activity was unrelated to the present work. The remaining authors declare no competing interests.

