## Supplementary Material for "Multiscale reorganization of brain and behavior under large-scale electrical perturbation"

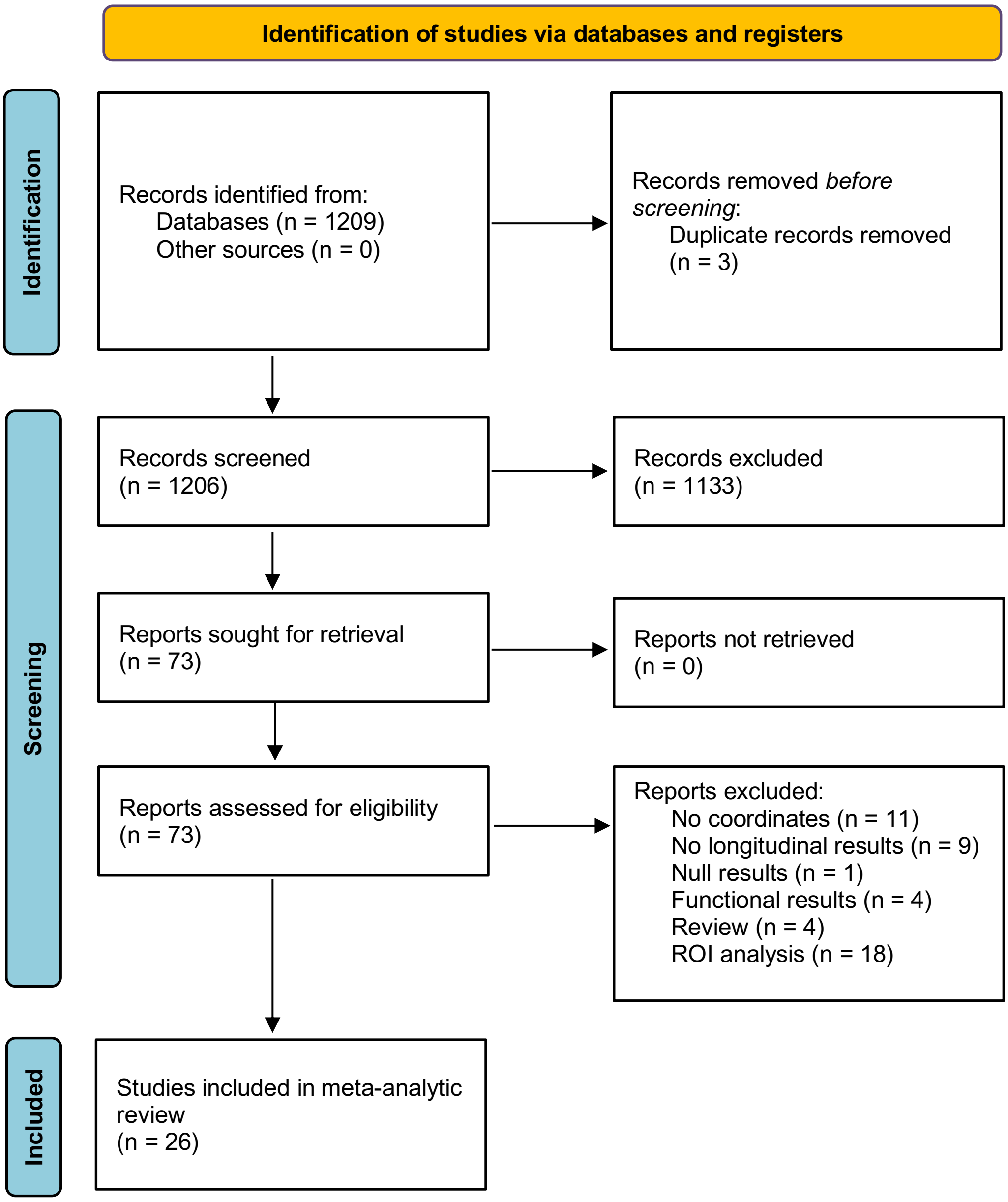


**Fig. S1 | PRISMA 2020 flow diagram of ECT-induced structural changes.** ECT, electroconvulsive therapy; ROI, region of interest.

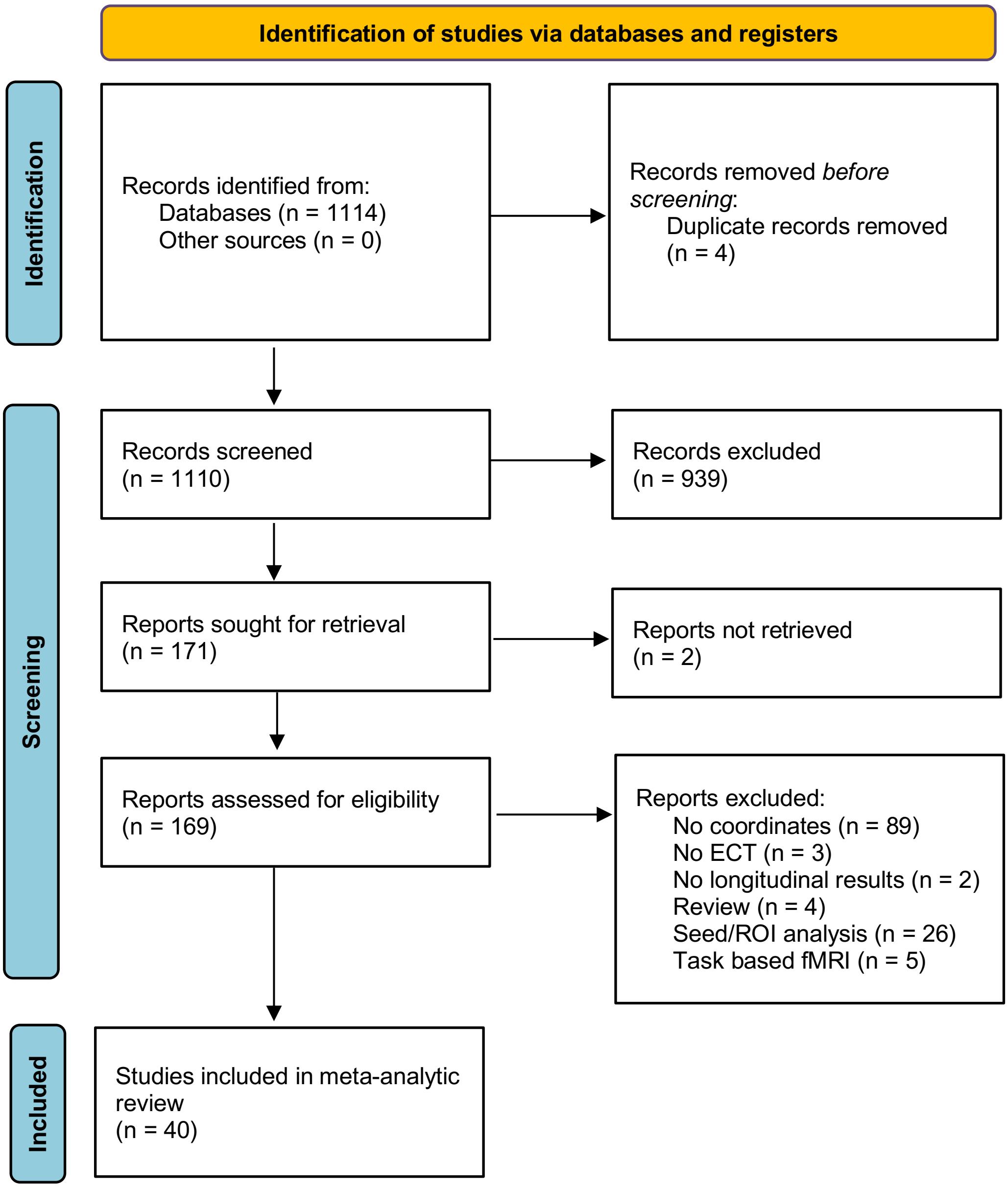


**Fig. S2 | PRISMA 2020 flow diagram of ECT-induced functional changes.** ECT, electroconvulsive therapy; ROI, region of interest.

| **Study** | **Subjects** | | | | **Imaging** | **Group-Level Analysis** | **Foci** | **Direction** |
|---|---|---|---|---|---|---|---|---|
| | ***ECT*** | | ***Comparison*** | | | | | |
| | *Disorder* | *N* | *Disorder* | *N* | | | | |
| **Borgers et al., 2024[1]** | MDD | 17 | MDD/HC | 33/21 | VBM | Group (ECT-MDD vs. MDD vs. HC) × Time (Before vs. After ECT vs. 2-year follow-up ECT) IA | 4 | |
| | MDD | 17 | | | VBM | Before < After ECT | 7 | + |
| | MDD | 17 | MDD/HC | 33/21 | VBM | Group (ECT-MDD vs. MDD vs. HC) × Time (Before vs. After ECT vs. 2-year follow-up ECT) IA (controlling for depression severity) | 4 | |
| | MDD | 17 | | | VBM | Before < After ECT (controlling for depression severity) | 5 | + |
| **Bouckaert et al., 2016[2]** | MDD | 28 | | | VBM | Before < After ECT | 9 | + |
| **Camilleri et al., 2020[3]** | MDD | 85 | | | VBM | Before < After ECT | 8 | + |
| **Cano et al., 2017[4]** | MDD | 12 | HC | 10 | VBM | Before < After ECT (after 9 sessions), HC < MDD | 7 | + |
| | MDD | 12 | HC | 10 | VBM | Before < After ECT (15 days after ECT completion), HC < MDD | 5 | + |
| **Cano et al., 2019[5]** | MDD | 12 | | | VBM | Before < After ECT, RUL < BT ECT | 2 | |
| **Cano et al., 2022[6]** | MDD | 15 | | | VBM | Before < After ECT | 4 | + |
| **Cano et al., 2023[7]** | MDD | 15 | | | VBM | Before < After ECT | 4 | + |
| | MDD | 15 | MDD | 17 | VBM | Before < After ECT, MDD-ECT > MDD-TMS | 3 | + |
| **Dukart et al., 2014[8]** | MDD/BPD | 10 | | | VBM | Before > After ECT | 1 | - |
| | MDD/BPD | 10 | | | VBM | Before < After ECT | 3 | + |
| **Gyger et al., 2021[9]** | MDD/BPD/SAD | 9 | | | VBM | Before vs. After ECT | 11 | |
| | MDD/BPD/SAD | 9 | | | VBM | Before ECT vs. Follow-up | 3 | |
| | MDD/BPD/SAD | 9 | | | VBM | MADRS × Time (Before ECT vs. After first ECT / After ECT / Follow-up) IA | 21 | |
| | MDD/BPD/SAD | 9 | | | VBM | MADRS × Time (Before ECT vs. Follow-up) IA | 30 | |
| **Gyger et al., 2021[10]** | MDD/BPD | 9 | | | VBM | Before < After ECT | 1 | + |
| **Kawashima et al., 2023[11]** | MDD | 18 | | | VBM | Before < After ECT | 39 | + |
| | SCZ | 12 | | | VBM | Before < After ECT | 12 | + |
| **Li et al., 2022[12]** | MDD | 30 | | | VBM | Before < After ECT | 2 | + |
| **Long et al., 2025[13]** | MDD | 29 | | | VBM | Before < After ECT | 18 | + |
| | MDD | 29 | KI/TSD | 43/55 | VBM | Group (ECT vs. KI vs. TSD) x Time (Before vs. After ECT) | 15 | |
| **Ota et al., 2015[14]** | MDD | 15 | | | VBM | Before < After ECT | 6 | + |
| **Qi et al., 2020[15]** | MDD | 118 | | | VBM | Before < After ECT | 21 | + |
| | MDD | 64 | | | VBM | Before < After ECT (responders) | 10 | + |
| | MDD | 54 | | | VBM | Before < After ECT (non-responders) | 2 | + |
| **Redlich et al., 2016[16]** | MDD | 23 | MED/HC | 23/21 | VBM | Group (ECT vs. MED vs. HC) × Time (Before vs. After ECT) IA | 1 | |
| | MDD | 23 | | | VBM | Before < After ECT | 4 | + |
| **Sartorius et al., 2016[17]** | MDD | 18 | | | VBM | Before < After ECT | 18 | + |
| **Sartorius et al., 2019[18]** | MDD | 92 | | | VBM | Before < After ECT | 6 | + |
| **Shan et al., 2021[19]** | SCZ | 27 | SCZ | 40 | VBM | Group (ECT-SCZ vs. Drug-SCZ) × Time (Before vs. After ECT) IA | 8 | |

| | | | | | | | | |
|---|---|---|---|---|---|---|---|---|
| | SCZ | 27 | | | VBM | Before < After ECT | 3 | + |
| | SCZ | 27 | | | VBM | Before < After ECT (TFCE corrected) | 1 | + |
| | SCZ | 27 | | | VBM | Before < After ECT (FWE corrected) | 3 | + |
| | SCZ | 16 | | | VBM | Before < After ECT (w/o patients on mood stabilizers) | 2 | + |
| **Takamiya et al., 2021**[20] | MDD/BPD | 27 | HC | 21 | VBM | Group (MDD vs. HC) × Time (Before vs. After ECT) IA | 18 | |
| | MDD/BPD | 20 | MDD/BPD | 7 | VBM | Group (remitters vs. nonremitters/controls) × Time (Before vs. After ECT) | 10 | |
| **Takamiya et al., 2022**[21] | MDD-P | 56 | | | VBM | Before < After ECT | 34 | + |
| | MDD-NP | 52 | | | VBM | Before < After ECT | 26 | + |
| **Thomann et al., 2017**[22] | MDD/SCZ | 21 | | | VBM | Before < After ECT | 3 | + |
| **van de Mortel et al., 2022**[23] | MDD/BPD | 88 | | | VBM | Before < After ECT | 10 | + |
| | MDD/BPD | 53 | | | VBM | Before < After ECT (Patients receiving RUL-ECT) | 16 | + |
| **Wang et al., 2017**[24] | MDD | 23 | | | VBM | Before < After ECT | 1 | + |
| **Wang et al., 2019**[25] | SCZ | 21 | SCZ | 21 | VBM | Group (SCZ-ECT vs. SCZ-Drug) × Time (Before vs. After ECT) IA | 4 | |
| **Xu et al., 2019**[26] | MDD | 11 | | | VBM | Correlation between ECT-induced change in global GMV and GMV | 16 | + |

**Table S1 | Studies reporting neuroimaging experiments indicating neuroplastic brain changes induced by electroconvulsive therapy.** BPD, bipolar disorder; BT, bitemporal; ECT, electroconvulsive therapy; FWE, familywise error; HC, healthy controls; IA, interaction; KI, ketamine infusion; MDD, major depressive disorder; MED, medication; RUL, right unilateral; SAD, schizoaffective disorder; SCZ, schizophrenia; TFCE, threshold-free cluster enhancement; TSD, total sleep deprivation; VBM, voxel-based morphometry; -NP, nonpsychotic; -P, psychotic.

| Study | Subjects | | | | Imaging | Group-Level Analysis | | Foci | Direction |
|---|---|---|---|---|---|---|---|---|---|
| | *ECT* | | *Comparison* | | | *Measure* | *Comparison* | | |
| | *Disorder* | *N* | *Disorder* | *N* | | | | | |
| **Argyelan et al., 2016[27]** | MDD/BPD | 13 | | | MRI | fALFF | Before > After ECT | 7 | - |
| **Bai et al., 2019[28]** | MDD/BPD | 28 | | | MRI | ALFF | Before < After ECT | 1 | + |
| | MDD/BPD | 33 | | | MRI | ALFF | Before > After ECT | 7 | - |
| | MDD/BPD | 33 | | | MRI | ALFF | Before < After ECT | 3 | + |
| **Belge et al., 2021[29]** | MDD/BPD | 17 | | | MRI | whole-brain FC of Anterior DMN | Before < After ECT | 1 | + |
| | MDD/BPD | 17 | | | MRI | whole-brain FC of Left CNN | Before < After ECT | 3 | + |
| | MDD/BPD | 17 | | | MRI | whole-brain FC of Right CNN | Before < After ECT | 2 | + |
| | MDD/BPD | 17 | | | MRI | whole-brain FC of SCN | Before < After ECT | 2 | + |
| **Du et al., 2016[30]** | MDD | 11 | | | MRI | ALFF | Before > After MECT | 3 | - |
| | MDD | 11 | | | MRI | ALFF | Before > After MECT (AlphaSim corrected) | 1 | - |
| | MDD | 11 | | | MRI | fALFF | Before > After MECT | 1 | - |
| **Fan et al., 2024[31]** | MDD | 42 | HC | 42 | MRI | brain entropy | Group (MDD vs. HC) x Time (Before vs. After ECT) IA | 2 | |
| | MDD | 42 | | | MRI | brain entropy | Before > After ECT | 2 | - |
| **Guo et al., 2024[32]** | MDD | 80 | HC | 37 | MRI | modular variability | Group (MDD vs. HC) x Time (Before vs. After ECT) IA | 29 | |
| | MDD | 80 | | | MRI | modular variability | Before > After ECT | 28 | - |
| | MDD-SI | 50 | | | MRI | modular variability | Before > After ECT | 1 | - |
| **Huang et al., 2018[33]** | SCZ | 21 | MED | 21 | MRI | global FC density | SCZ group (MECT + Drug vs. Drug) x Time (Before vs. After MECT) | 3 | |
| | SCZ | 21 | | | MRI | global FC density | Before < After ECT | 3 | + |
| | SCZ | 21 | | | MRI | global FC density | Before < After ECT ($p<0.005$) | 3 | + |
| | SCZ | 21 | | | MRI | global FC density | Before < After ECT | 7 | + |
| | SCZ | 21 | | | MRI | global FC density | Before > After ECT | 5 | - |
| **Kohn et al., 2007[34]** | MDD | 7 | MED | 17 | SPECT | rCBF | MDD group (Drug- vs. ECT-responders) x Time (After vs. Before ECT) | 1 | |
| | MDD | 7 | | | SPECT | rCBF | Before > After ECT (responders) | 1 | - |
| **Kong et al., 2017[35]** | MDD | 13 | | | MRI | ReHo | Before > After ECT | 2 | - |
| | MDD | 13 | | | MRI | ALFF | Before > After ECT | 3 | - |
| | MDD | 13 | | | MRI | ALFF | Before < After ECT | 2 | + |
| **Leaver et al., 2016[36]** | MDD/BPD | 30 | | | MRI | resting-state FC within Th/vBGN | Before < After ECT | 1 | + |
| | MDD/BPD | 30 | | | MRI | resting-state FC within Anterior DMN | Before < After ECT | 1 | + |
| | MDD/BPD | 30 | | | MRI | resting-state FC within SN | Before < After ECT | 1 | + |
| | MDD/BPD | 30 | | | MRI | resting-state FC within SN | Before > After ECT | 1 | - |
| | MDD/BPD | 30 | | | MRI | resting-state FC within Posterior DMN | Before < After ECT | 1 | + |
| | MDD/BPD | 30 | | | MRI | resting-state FC within AMTN | Before > After ECT | 1 | - |
| **Leaver et al., 2019[37]** | MDD/BPD | 51 | | | ASL-MRI | rCBF | Before < After 2 ECT sessions | 1 | + |
| | MDD/BPD | 51 | | | ASL-MRI | rCBF | Before > After 2 ECT sessions | 1 | - |

| | | | | | | | |
|---|---|---|---|---|---|---|---|
| | MDD/BPD | 43 | ASL-MRI | rCBF | Before < After ECT | 3 | + |
| | MDD/BPD | 17 | ASL-MRI | rCBF | Before < After ECT (responders) | 3 | + |
| | MDD/BPD | 17 | ASL-MRI | rCBF | Before > After ECT (responders) | 3 | - |
| | MDD/BPD | 26 | ASL-MRI | rCBF | Before < After ECT (non-responders) | 1 | + |
| | MDD/BPD | 17 | ASL-MRI | rCBF | Before > After ECT (non-responders) | 1 | - |
| **Li et al., 2019[38]** | MDD | 24 | MRI | global FC density | Before < After MECT | 3 | + |
| | MDD | 24 | MRI | global FC density | Before > After MECT | 1 | - |
| **Li et al., 2021[39]** | MDD-SI | 14 | MRI | ALFF | Before > After ECT | 1 | - |
| | MDD-SI | 14 | MRI | fALFF | Before > After ECT | 1 | - |
| | MDD-SI | 14 | MRI | degree centrality | Before > After ECT | 1 | - |
| | MDD-SI | 14 | MRI | degree centrality | Before < After ECT | 1 | + |
| **Li et al., 2022[40]** | MDD-SI | 30 | MRI | ALFF | Before < After ECT | 1 | + |
| | MDD-SI | 30 | MRI | ALFF | Before > After ECT | 3 | - |
| | MDD-SI | 30 | MRI | ReHo | Before < After ECT | 2 | + |
| **Li et al., 2022[41]** | MDD | 22 | MRI | ALFF | Before < After ECT | 5 | + |
| | MDD | 22 | MRI | ALFF | Before > After ECT | 1 | - |
| | MDD | 22 | MRI | ALFF (slow-5) | Before < After ECT | 5 | + |
| | MDD | 22 | MRI | ALFF (slow-5) | Before > After ECT | 3 | - |
| | MDD | 22 | MRI | ALFF (slow-4) | Before < After ECT | 4 | + |
| **Liu et al., 2015[42]** | MDD | 23 | MRI | ALFF | Before < After ECT | 6 | + |
| | MDD | 23 | MRI | ALFF | Before > After ECT | 1 | - |
| **Liu et al., 2022[43]** | MDD | 9 | MRI | nodal degree | Before < After ECT | 2 | + |
| | MDD | 9 | MRI | nodal degree | Before > After ECT | 4 | - |
| | MDD | 9 | MRI | nodal flexibility | Before < After ECT | 1 | + |
| | MDD | 9 | MRI | nodal flexibility | Before > After ECT | 18 | - |
| **McCormick et al., 2007[44]** | MDD-P | 10 | PET | regional glucose rate | Before < After ECT | 1 | + |
| | MDD-P | 10 | PET | regional glucose rate | Before > After ECT | 1 | - |
| | MDD-P | 10 | PET | rCMRGlu | Association with HAM-D score changes, Before < After ECT | 2 | + |
| | MDD-P | 10 | PET | rCMRGlu | Association with SAPS score changes, Before < After ECT | 2 | + |
| **Mo et al., 2020[45]** | MDD | 28 | MRI | ReHo | Before < After ECT | 1 | + |
| **Mulders et al., 2016[46]** | MDD | 16 | MRI | DMN coherence | Before > After ECT | 1 | - |
| | MDD | 8 | MRI | DMN coherence | Before > After ECT (responders) | 1 | - |
| | MDD | 8 | MRI | DMN coherence | Before < After ECT (responders) | 2 | + |
| | MDD | 8 | MRI | DMN coherence | Before > After ECT (non-responders) | 2 | - |
| **Nie et al., 2022[47]** | MDD/BPD | 45 | MRI | dALFF | Before < After ECT | 1 | + |
| | MDD/BPD | 45 | MRI | dALFF | Before > After ECT | 2 | - |
| **Nobler et al., 2001[48]** | MDD/BPD | 10 | PET | rCMRGlu | Before > After ECT | 4 | - |

| | | | | | | | |
|---|---|---|---|---|---|---|---|
| **Pang et al., 2022**[49] | MDD | 30 | MRI | FC (multi-voxel pattern analysis) | Before vs. After ECT | 13 | |
| **Perrin et al., 2012**[50] | MDD | 9 | MRI | average global FC | Before > After ECT | 4 | - |
| **Qi et al., 2020**[15] | MDD | 118 | MRI | fALFF in responsive joint components | Before > After ECT | 13 | - |
| | MDD | 64 | MRI | fALFF in responsive joint components | Before > After ECT (responders) | 8 | - |
| | MDD | 54 | MRI | fALFF in responsive joint components | Before > After ECT (non-responders) | 9 | - |
| **Qiu et al., 2019**[51] | MDD | 24 | MRI | fALFF | Before > After ECT | 3 | - |
| **Segawa et al., 2006**[52] | MDD/BPD | 10 | SPECT | rCBF | Negative correlation with HAM-D scores | 7 | - |
| **Shi et al., 2022**[53] | MDD | 10 | ASL-MRI | rCBF | Before > After ECT | 2 | - |
| **Suwa et al., 2012**[54] | MDD/BPD | 16 | PET | regional glucose metabolism | Before > After ECT | 8 | - |
| | MDD/BPD | 16 | PET | regional glucose metabolism | Before < After ECT | 2 | + |
| | MDD/BPD | 12 | PET | regional glucose metabolism | Before > After ECT (responders) | 8 | - |
| | MDD/BPD | 12 | PET | regional glucose metabolism | Before < After ECT (responders) | 2 | + |
| **Takamiya et al., 2021**[55] | MDD/BPD | 27 | MRI | FC (multi-voxel pattern analysis) | Association with HAM-D score changes | 6 | |
| | MDD/BPD | 27 | MRI | FC (multi-voxel pattern analysis) | Association with MMSE score changes | 2 | |
| **Tong et al., 2022**[56] | SCZ | 29 | MRI | ReHo | Before < After 1 ECT session | 3 | + |
| | SCZ | 27 | MRI | ReHo | Before < After ECT | 4 | + |
| | SCZ | 27 | MRI | ReHo | After 1 ECT session < After ECT | 2 | + |
| | SCZ | 27 | MRI | ReHo | After 1 ECT session > After ECT | 2 | - |
| **Usui et al., 2011**[57] | PDP | 8 | SPECT | rCBF | Before < After ECT | 8 | + |
| | PDP | 8 | SPECT | rCBF | Before < After ECT (corrected for multiple comparison) | 1 | + |
| | PDP | 8 | SPECT | rCBF | Before > After ECT | 4 | - |
| **Wang et al., 2018**[58] | MDD | 23 | MRI | local FC density | Negative correlation with HAM-D scores | 3 | + |
| **Wang et al., 2020**[59] | MDD | 23 | MRI | FC homogeneity | Before < After ECT | 2 | + |
| **Wang et al., 2023**[60] | MDD-SI | 23 | MRI | fALFF | Before < After ECT (responders) | 1 | + |
| **Wang et al., 2024**[61] | MDD | 32 | MRI | degree centrality | Before < After ECT | 1 | + |
| | MDD | 32 | MRI | degree centrality | Before > After ECT | 1 | - |
| **Wei et al., 2014**[62] | MDD | 11 | MRI | voxel-mirrored homotopic connectivity | Before < After ECT | 8 | + |
| **Wei et al., 2018**[63] | MDD | 26 | MRI | FC strength | Before < After ECT | 1 | + |
| **Yuuki et al., 2005**[64] | MDD/BPD | 7 | PET | rCMRGlu | Before > After ECT | 1 | - |
| | MDD/BPD | 7 | PET | rCMRGlu | Before < After ECT | 2 | + |
| **Zhang et al., 2021**[65] | MDD | 46 | MRI | ALFF | Before < After ECT | 3 | + |
| | MDD | 46 | MRI | degree centrality | Before < After ECT | 9 | + |

**Table S2 | Studies reporting neuroimaging experiments indicating functional brain changes induced by electroconvulsive therapy.** ALFF, amplitude of low frequency fluctuation; AMTN, anteromedial temporal network; ASL, arterial spin-labeled; BPD, bipolar disorder; CEN, cognitive executive network; dALFF, dynamic amplitude of low-frequency fluctuation; DMN, default mode network; ECT, electroconvulsive therapy; fALFF, fractional amplitude of low frequency fluctuation; HAM-D, Hamilton Depression Rating Scale; HC, healthy controls; IA, interaction; MDD, major depressive disorder; MECT, modified electroconvulsive therapy; MED, medication; MMSE, mini-mental state examination; MRI, magnetic resonance imaging; P, psychotic; PDP, Parkinson's disease psychosis; PET, positron emission tomography; rCBF, regional cerebral blood flow; rCMRGlu, regional cerebral metabolism rate of glucose; ReHo, regional homogeneity; SAPS, Scale for the Assessment of Positive Symptoms; SCN, subcortical network; SCZ, schizophrenia; SN, salience network; SPECT, single photon emission computed tomography; Th/vBGN, thalamus and ventral basal ganglia network; -NSI, no suicidal ideation; -SI, suicidal ideation.

| | Macroanatomical Location | Cytoarchitectonic Location | Cluster Size in Voxels | MNI Coordinates | | | TFCE Score |
|---|---|---|---|---|---|---|---|
| | | | | x | y | z | |
| R | Amygdala/Hippocampus | VTM/CA1/LB | 2992 | 26 | -8 | -20 | $6.24 \times 10^4$ |
| R | Amygdala | LB | | 32 | 2 | -30 | $2.89 \times 10^4$ |
| R | Temporal pole | | | 36 | 14 | -28 | $2.63 \times 10^4$ |
| R | Temporal pole | | | 38 | 10 | -22 | $2.62 \times 10^4$ |
| R | Central opercular cortex | Id4/OP3 | | 38 | 0 | 16 | $1.90 \times 10^4$ |
| R | Insular cortex | Id5/Ia | | 44 | -2 | 0 | $1.56 \times 10^4$ |
| R | Insular cortex | Id5 | | 42 | -2 | 0 | $1.56 \times 10^4$ |
| R | Planum polare / Insular cortex | Id3/Ia | | 42 | 0 | -14 | $1.45 \times 10^4$ |
| R | Planum polare / Central opercular cortex | TE 1.2 / OP4 | | 54 | 4 | -2 | $1.24 \times 10^4$ |
| R | Superior temporal cortex / Insular cortex | TE 4 | | 46 | -16 | -10 | $1.07 \times 10^4$ |
| L | Amygdala | MF/CM/SF | 725 | -18 | -8 | -14 | $2.01 \times 10^4$ |
| L | Hippocampus | DG/CA1/Subiculum | | -28 | -18 | -20 | $1.39 \times 10^4$ |
| L | Hippocampus | DG/CA1/Subiculum | | -28 | -16 | -20 | $1.39 \times 10^4$ |
| L | Hippocampus | | | -32 | -24 | -8 | $8.97 \times 10^3$ |
| L | Putamen | | | -30 | -20 | -6 | $8.88 \times 10^3$ |
| L | Ventral striatum | | | -22 | 8 | -10 | $7.99 \times 10^3$ |
| | Subgenual cingulate cortex | 25/33 | 221 | 0 | 8 | -6 | $1.08 \times 10^4$ |
| R | Ventral striatum | | | 6 | 12 | 0 | $1.07 \times 10^4$ |
| R | Ventral striatum | s24 | 10 | 8 | 20 | -8 | $7.41 \times 10^3$ |

**Table S3 | Electroconvulsive therapy-induced changes in brain morphology.** Across 45 experiments featuring 408 foci of changes in gray matter induced by electroconvulsive therapy, ALE revealed four clusters of significant convergence. Results are corrected for multiple comparisons using threshold-free cluster enhancement ($p<0.05$). ALE, activation likelihood estimation; MNI, Montreal Neurological Institute.

| | Macroanatomical Location | Cytoarchitectonic Location | Cluster Size in Voxels | MNI Coordinates | | | TFCE Score |
|---|---|---|---|---|---|---|---|
| | | | | x | y | z | |
| L | Temporoparietal junction | PGp/PGa | 118 | -42 | -68 | 40 | $1.19 \times 10^4$ |
| L | Temporoparietal junction | PGp/PGa | | -46 | -62 | 30 | $1.04 \times 10^4$ |
| L | Temporoparietal junction | PGa/PFm | | -46 | -60 | 30 | $1.04 \times 10^4$ |
| L | Temporoparietal junction | PGa/PFm | | -44 | -62 | 32 | $1.04 \times 10^4$ |
| L | Temporoparietal junction | PFm/PGa/hIP1 | | -46 | -56 | 38 | $9.26 \times 10^3$ |
| R | Insular cortex | Ia | 25 | 42 | 6 | -8 | $1.06 \times 10^4$ |
| | Anterior midcingulate cortex | 33 | 25 | -6 | 14 | 28 | $1.20 \times 10^4$ |
| | Medial frontal pole | | 23 | 0 | 60 | 18 | $9.93 \times 10^3$ |
| R | Temporoparietal junction | PGp/PGa | 23 | 48 | -62 | 34 | $1.02 \times 10^4$ |
| | Posterior cingulate cortex | | 22 | 4 | -38 | 28 | $1.02 \times 10^4$ |
| L | Middle temporal gyrus | | 20 | -60 | -28 | -16 | $1.08 \times 10^4$ |
| L | Medial frontal pole | Fp1/Fp2 | 18 | -12 | 66 | 4 | $9.57 \times 10^3$ |
| L | Medial frontal pole | Fp2/p32 | | -6 | 60 | 6 | $9.56 \times 10^4$ |
| R | Hippocampus | CA1/Subiculum | 16 | 32 | -16 | -24 | $9.59 \times 10^3$ |
| R | Medial frontal pole | Fp2/p32 | 7 | 6 | 62 | 8 | $9.40 \times 10^3$ |

**Table S4 | Electroconvulsive therapy-induced changes in brain physiology.** Across 103 experiments featuring 340 foci of physiological changes induced by electroconvulsive therapy, ALE revealed ten clusters of significant convergence. Results are corrected for multiple comparisons using threshold-free cluster enhancement ($p<0.05$). ALE, activation likelihood estimation; MNI, Montreal Neurological Institute.

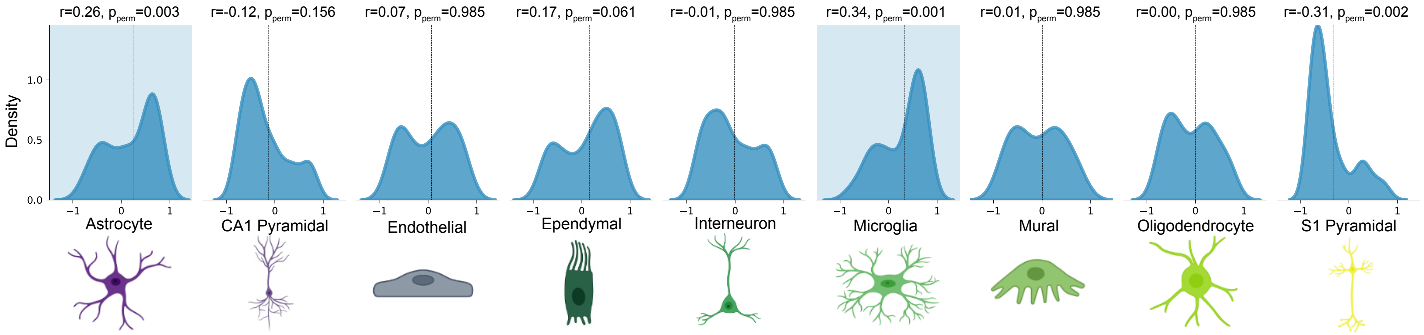


**Fig. S3 | Cell-type association of ECT-induced structural changes in the right hemisphere.** The virtual histology approach was applied to the right hemisphere. Identical to Fig. 1b, the x-axis indicates the Spearman's rank correlation coefficient between ECT-induced structural changes and the expression profiles of a given set of cell-type specific genes. The y-axis represents the estimated probability density of the correlation coefficients. The vertical dashed line marks the average correlation coefficient across all genes specific to the cell type shown in each panel. A colored background highlights positive average correlations with $p_{perm} \leq 0.05$, corrected for multiple comparisons using FWER correction.

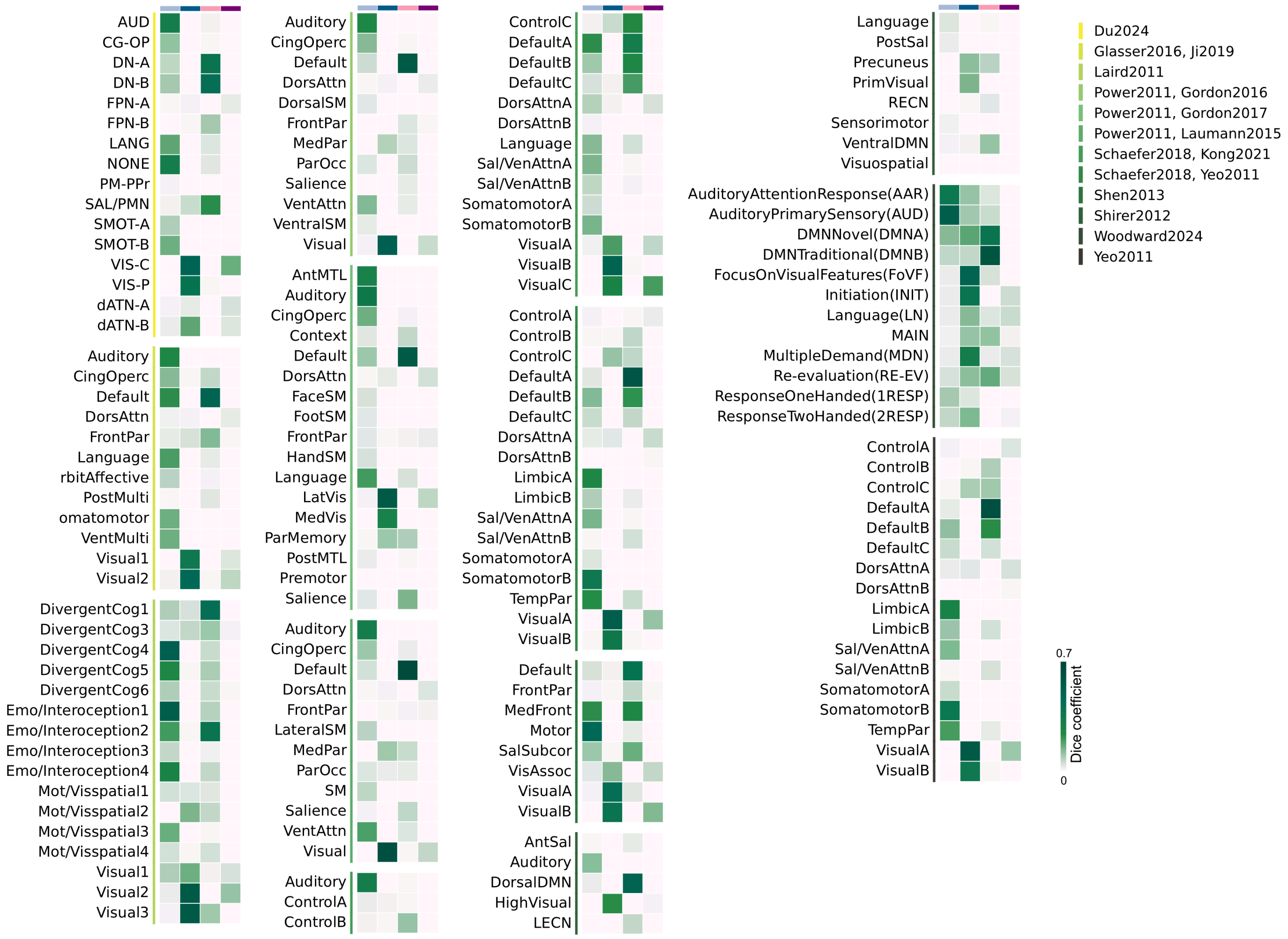


**Fig. S4 | Network correlates of ECT.** Each ECT network map was analyzed with the Network Correspondence Toolbox. Dice coefficients are shown in the heat map for all networks, sorted by atlas (top right), using the same color scheme as before: Positive structural network (light blue), negative structural network (dark blue), positive functional network (light pink), and negative functional network (purple).

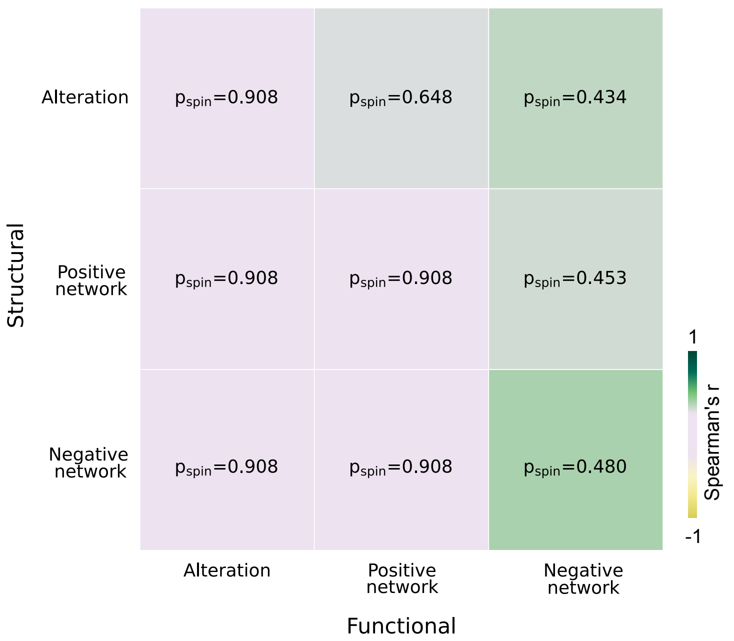


**Fig. S5 | Comparison between structural and functional maps.** Spearman's rank correlation was calculated between all structural and functional maps. Significance was assessed using spatial autocorrelation-preserving null models (spin tests) with family-wise error rate (FWE) correction for multiple comparisons.

| Receptor/ transporter | Neurotransmitter | Tracer | Measure | N (males) | Age | References |
|---|---|---|---|---|---|---|
| **5-HT$_{1A}$** | Serotonin | [$^{11}$C]WAY-100635 | BP$_{ND}$ | 35(18) | 26.3±5.2 | Savli et al. 2012[66] |
| **5-HT$_{1B}$** | Serotonin | [$^{11}$C]943 | BP$_{ND}$ | 65(16) | 33.7±9.7 | Gallezot et al. 2010[67] |
| **5-HT$_{1B}$** | Serotonin | [$^{11}$C]943 | BP$_{ND}$ | 23(15) | 28.7±7 | Savli et al. 2012[66] |
| **5-HT$_{2A}$** | Serotonin | [$^{18}$F]Altanserin | BP$_{ND}$ | 19(11) | 28.2±5.7 | Savli et al. 2012[66] |
| **5-HT$_{4}$** | Serotonin | [$^{11}$C]SB207145 | B$_{max}$ | 59(41) | 25.9±5.3 | Beliveau et al. 2017[68] |
| **5-HT$_{6}$** | Serotonin | [$^{11}$C]GSK215083 | BP$_{ND}$ | 30(30) | 36.6±9.0 | Radhakrishnan et al. 2020[69] |
| **5-HTT** | Serotonin | [$^{11}$C]DASB | BP$_{ND}$ | 18(12) | 30.5±9.5 | Savli et al. 2012[66] |
| **D$_{1}$** | Dopamine | [$^{11}$C]233390 | BP$_{ND}$ | 13(6) | 33 ±13 | Kaller et al. 2017[70] |
| **D$_{2}$** | Dopamine | [$^{11}$C]FLB-457 | BP$_{ND}$ | 37(17) | 48.4 ± 16.9 | Smith et al. 2019[71] |
| **D$_{2}$** | Dopamine | [$^{11}$C]FLB-457 | BP$_{ND}$ | 55(26) | 32.5 ± 9.7 | Sandiego et al. 2015[72] |
| **DAT$^{T}$** | Dopamine | [$^{123}$I]-FP-CIT | SUVR | 174(109) | 61±11 | Dukart et al. 2018[73] |
| **GABA$_{A}$** | GABA | [$^{11}$C]Flumazenil | BP$_{ND}$ | 6(6) | 43±4 | Dukart et al. 2018[73] |
| **mGluR$_{5}$** | Glutamate | [$^{11}$C]ABP688 | BP$_{ND}$ | 28(15) | 33.1±11.2 | DuBois et al. 2016[74] |
| **mGluR$_{5}$** | Glutamate | [$^{11}$C]ABP688 | BP$_{ND}$ | 22(12) | 67.9±9.6 | Hansen et al. 2022[75] |
| **mGluR$_{5}$** | Glutamate | [$^{11}$C]ABP688 | BP$_{ND}$ | 73(25) | 29.9±3.0 | Smart et al. 2019[76] |
| **H$_{3}$** | Histamine | [$^{11}$C]GSK189254 | BP$_{ND}$ | 8(7) | 31.7±9.0 | Gallezot et al. 2017[77] |
| **NET$^{T}$** | Norepinephrine | [$^{11}$C]MRB | BP$_{ND}$ | 77(50) | 33.4±9.2 | Ding et al. 2010[78] |
| **MOR** | Opioid | [$^{11}$C]Carfentanil | BP$_{ND}$ | 204(132) | 32.3±10.8 | Kantonen et al. 2020[79] |
| **$\alpha_4\beta_2$** | Acetylcholine | [$^{18}$F]Flubatine | V$_{T}$ | 30(20) | 33.5±10.7 | Hillmer et al. 2016[80] |
| **M$_{1}$** | Acetylcholine | [$^{11}$C]LSN3172176 | BP$_{ND}$ | 24(13) | 40.5±11.7 | Naganawa et al. 2021[81] |
| **VAChT$^{T}$** | Acetylcholine | [$^{18}$F]FEOBV | SUVR | 18(5) | 66.8 ± 6.8 | Aghourian et al. 2017[82] |
| **VAChT$^{T}$** | Acetylcholine | [$^{18}$F]FEOBV | SUVR | 5(4) | 68.3 ± 3.1 | Bedard et al. 2019[83] |
| **VAChT$^{T}$** | Acetylcholine | [$^{18}$F]FEOBV | SUVR | 4(3) | 37 ± 10.2 | Hansen et al. 2022[75] |

**Table S5 | Neurotransmitter receptors and transporters.** BPND, non-displaceable binding potential; Bmax, converted from binding potential using autoradiography-derived densities; SUVR, standard uptake value ratio; VT, tracer distribution volume. The superscript T indicates transporters.

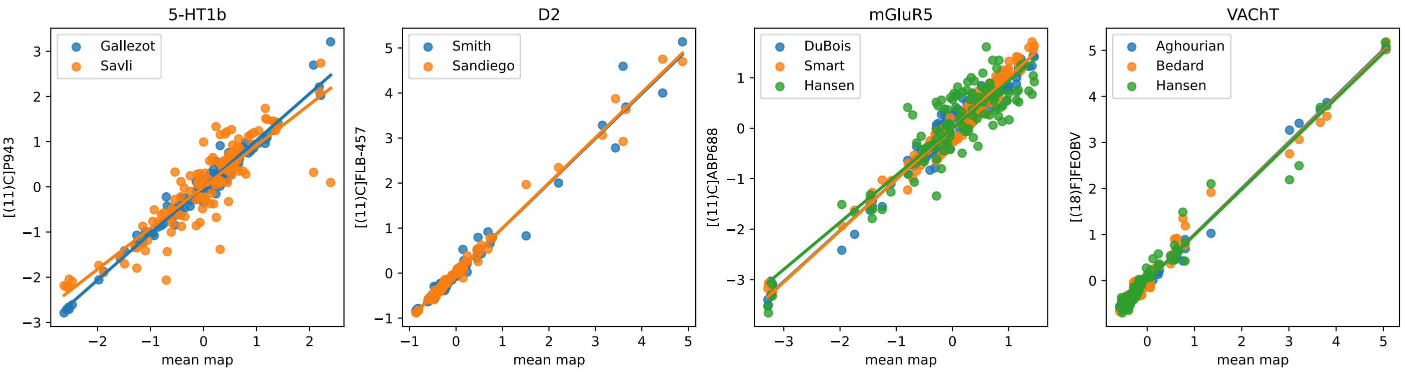


**Fig. S6 | Correlation of Neurotransmitter receptor and transporter densities.** Neurotransmitter receptor and transporter maps from the same tracer were combined into a single average neurotransmitter map. Each individual map (y-axis) is highly correlated with the mean map (x-axis), where names indicate the source of each individual map; see Supplementary Table S5.

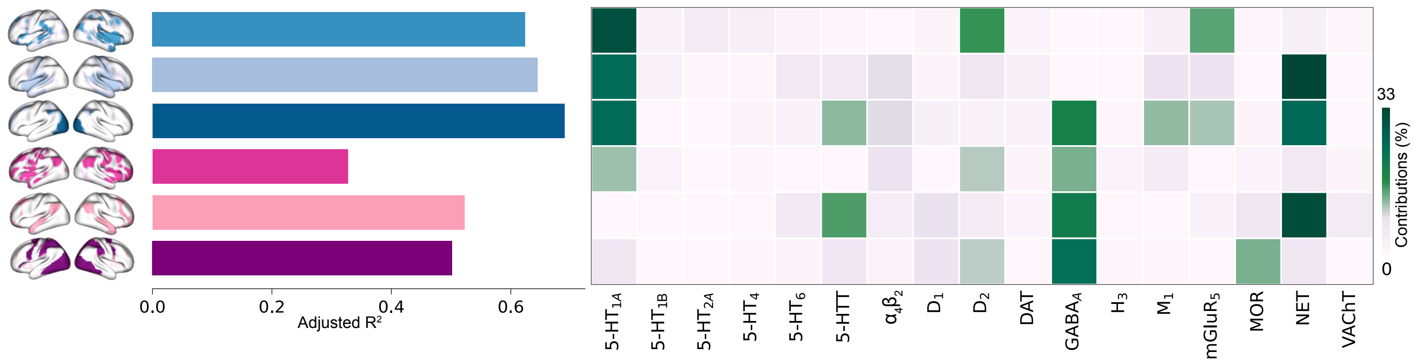


**Fig. S7 | Interactional dominance**. For each ECT map, a multiple linear regression model was fitted to predict its spatial profile based on 17 neurotransmitter receptor and transporter density maps. The ECT maps are shown on the left, and the thresholded maps are displayed in Fig. 1c, using the same color scheme: structural change (mid blue), positive structural network (light blue), negative structural network (dark blue), functional changes (pink), positive functional network (light pink), and negative functional network (purple). Model fits (adjusted $R^2$) are displayed in the bar plots identical to Fig. 4a. The heatmap shows interactional dominance, defined as the change in $R^2$ when an independent variable is added to the submodel where all other independent variables are present, normalized by the total $R^2$ of the model (one model per row).

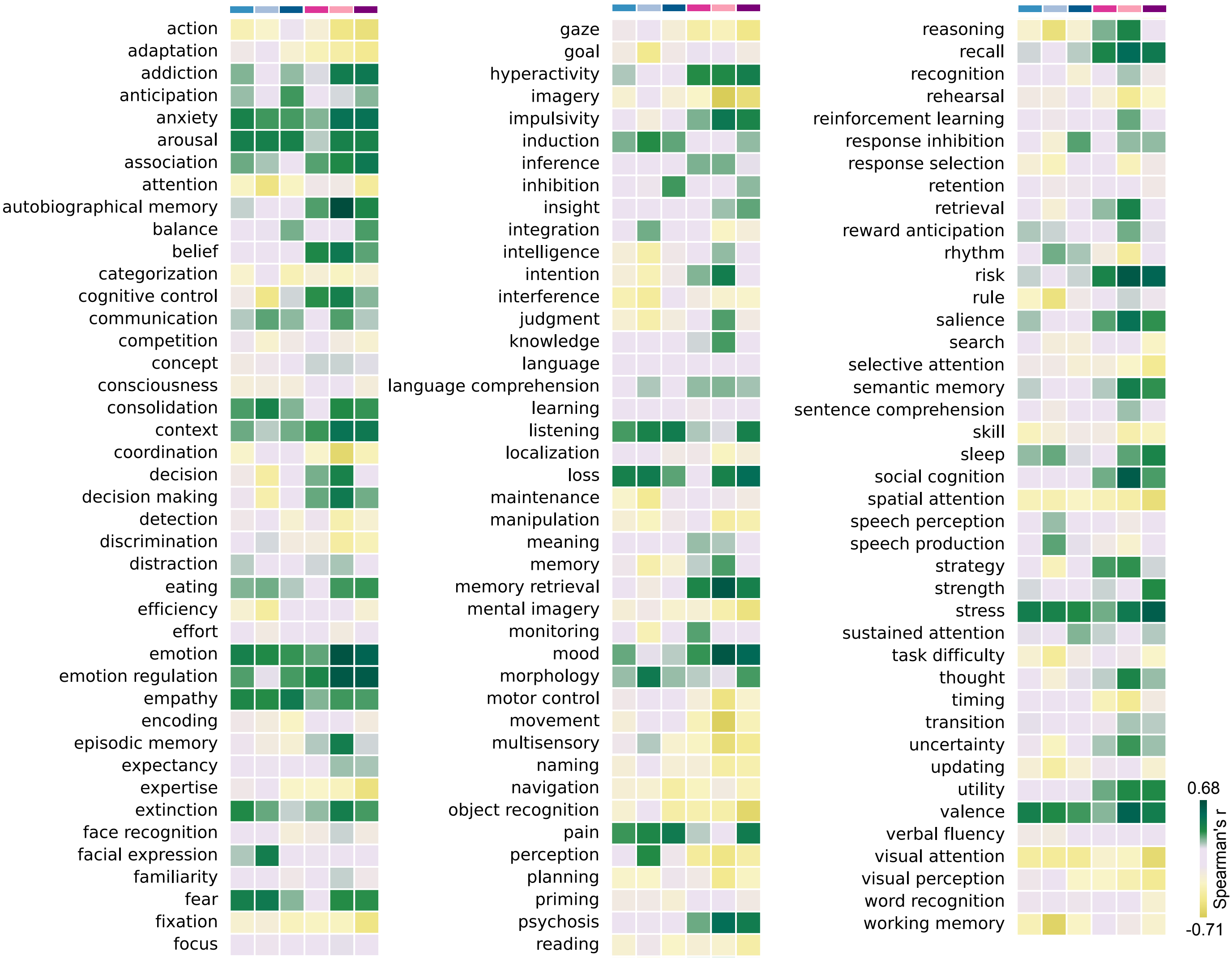


**Fig. S8 | Functional characterization of multimodal ECT effects.** Each ECT map was correlated with cognitive and behavioral meta-analytic activation maps from Neurosynth. Spearman's rank correlations are shown in the heat map for all 125 terms, using the same color scheme for the ECT maps as before: structural change (mid blue), positive structural network (light blue), negative structural network (dark blue), functional changes (pink), positive functional network (light pink), and negative functional network (purple).

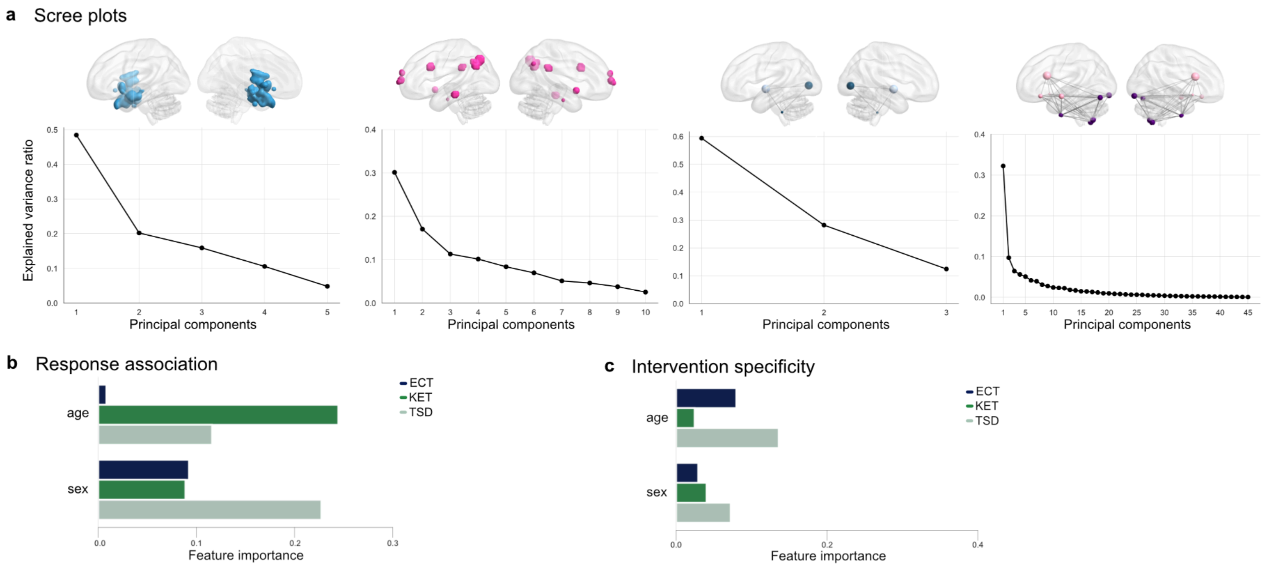


**Fig. S9 | Principal component analysis, response association, and ECT specificity a,** PCA was conducted separately for each ECT feature domain across all intervention groups in the fast-acting antidepressant dataset. Scree plots show principal components (x-axis) and explained variance ratio (y-axis), representing the proportion of total variance explained by each component. From left to right: changes in gray matter (structural ROIs), fractional amplitude of low-frequency fluctuations (fALFF; functional ROIs), structural network connectivity, and functional network connectivity. Elbow inspection suggested retaining the first 1, 2, 2, and 2 components, respectively. To ensure consistent dimensionality across domains and comparable influence of each domain in subsequent analyses, the mode (two principal components) was retained for all feature domains. **b,** and **c,** Feature importance of demographic covariates, normalized across all predictors. Domain feature importances were balanced prior to normalization.

| **Intervention** | **PC1 – SC** | **PC2 - SC** | **PC1 - FC** | **PC2 - FC** | **PC1 - SN** | **PC2 - SN** | **PC1 - FN** | **PC2 -FN** | **Age** | **Sex** |
|---|---|---|---|---|---|---|---|---|---|---|
| **ECT** | 0.141 | -0.455 | 0.488 | -1.579 | 0.807 | -1.307 | 0.810 | -0.649 | 0.0429 | -0.483 |
| **KET** | 0.553 | 0.016 | -0.047 | 0.211 | 0.093 | -0.392 | 0.566 | -0.519 | 0.711 | -0.258 |
| **TSD** | 0.027 | -0.053 | 0.359 | -0.333 | 0.587 | 0.165 | -0.229 | -0.348 | -0.278 | 0.545 |

**Table S6 | Ridge regression coefficients.** Raw coefficients prior to domain balancing and normalization from ridge regression models predicting clinical outcome (residualized QIDS change) for each intervention group. FC, functional connectivity; FN, functional network; KET, ketamine; PC1, first principal component; PC2, second principal component; SC, structural connectivity; SN, structural network; TSD, total sleep deprivation.

| **Intervention** | **PC1 – SC** | **PC2 - SC** | **PC1 - FC** | **PC2 - FC** | **PC1 - SN** | **PC2 - SN** | **PC1 - FN** | **PC2 -FN** | **Age** | **Sex** |
|---|---|---|---|---|---|---|---|---|---|---|
| **ECT** | -2.123 | 0.272 | -0.579 | -0.196 | -0.570 | -0.646 | 0.586 | -0.475 | 0.391 | -0.143 |
| **KET** | 1.041 | -0.025 | 0.634 | 0.332 | 0.121 | 0.230 | 0.098 | 0.184 | 0.058 | -0.096 |
| **TSD** | 1.088 | -0.247 | -0.056 | -0.136 | 0.449 | 0.416 | -0.683 | 0.291 | -0.450 | 0.239 |

**Table S7 | Multinomial logistic regression coefficients.** Raw coefficients for each intervention group, derived from a multinomial logistic regression predicting intervention group membership, prior to domain balancing and normalization. FC, functional connectivity; FN, functional network; KET, ketamine; PC1, first principal component; PC2, second principal component; SC, structural connectivity; SN, structural network; TSD, total sleep deprivation.
